# Indole produced during dysbiosis mediates host–microorganism chemical communication

**DOI:** 10.1101/2022.12.19.520989

**Authors:** Rui-Qiu Yang, Yong-Hong Chen, Qin-Yi Wu, Jie Tang, Shan-Zhuang Niu, Qiu Zhao, Yi-Cheng Ma, Cheng-Gang Zou

## Abstract

An imbalance of the gut microbiota, termed dysbiosis, has a substantial impact on host physiology. However, the mechanism by which host deals with gut dysbiosis to maintain fitness remains largely unknown. In *C. elegans*, *E. coli*, which is its bacterial diets, proliferates in its intestinal lumen during aging. Here, we demonstrate that progressive intestinal proliferation of *E. coli* activates the transcription factor DAF-16, which is required for maintenance of longevity and organismal fitness in worms with age. DAF-16 up-regulates two lysozymes *lys-7* and *lys-8*, thus limiting the bacterial accumulation in the gut of worms during aging. During dysbiosis, the levels of indole produced by *E. coli* are increased in worms. Indole is involved in the activation of DAF-16 by TRPA-1 in neurons of worms. Our finding demonstrates that indole functions as a microbial signal of gut dysbiosis to promote fitness of the host.

## Introduction

The microbiota in the gut has a substantial impact on host nutrition, metabolism, immune function, development, behavior, and lifespan (Lee & Brey 2013; Johnson & Foster 2018; Bana & Cabreiro 2019). Microbial community disequilibria, so-called dysbiosis, has been implicated in a broad range of human diseases, such as obesity, insulin resistance, autoimmune disorders, inflammatory bowel disease (IBD), aging and increased pathogen susceptibility (Honda & Littman 2012; Wu *et al*. 2015). Therefore, understanding of the mechanisms that modulate host–microbe interactions will provide important insight into treating these diseases by intervening the microbial communities.

The genetically tractable model organism *Caenorhabditis elegans* has contributed greatly to understand the role of host-microbiota interactions in host physiology (Cabreiro & Gems 2013; Zhang *et al*. 2017). In the bacterivore nematode, the microbiota in the gut can be easily manipulated, making it an excellent model for studying how the microbiota affect host physiology in the context of disease and aging at a single species (Cabreiro & Gems 2013). For instance, the neurotransmitter tyramine produced by intestinal Providencia bacteria can direct sensory behavioural decisions by modulate multiple monoaminergic pathways in *C. elegans* (O’Donnell *et al*. 2020). On the other hand, *C. elegans*-based studies have revealed a variety of signaling cascades involved in the innate immune responses against microbial infection (Irazoqui *et al*. 2010b), including the p38 mitogen activated protein kinase (MAPK)/PMK-1, ERK MAPK/MPK-1, the heat shock transcription factor HSF-1, the transforming growth factor (TGF)β/bone morphogenetic protein (BMP) signaling (Zugasti & Ewbank 2009), and the forkhead transcription factor DAF-16/FOXO pathway (Kim *et al*. 2002; Garsin *et al*. 2003; Singh & Aballay 2006; Zou *et al*. 2013). In Drosophila and mammalian cells, activation of FOXOs up-regulates a set of antimicrobial peptides, such as drosomycin and defensins (Becker *et al*. 2010), implicating that the role for FOXOs in innate immunity is conserved across species. Furthermore, disruption of these innate immune-reltaed pathways, such as the p38 MAPK pathway and the TGFβ/BMP signaling cascade, turns beneficial bacteria commensal to pathogenic in worms (Montalvo-Katz *et al*. 2013; Berg *et al*. 2019). These results indicate that the integrity of immune system is also essential for host defense against non-pathogenic bacteria.

*Escherichia coli* (strain OP50) is conventionally used as a bacterial food for culturing *C. elegans*. In general, most of *E. coli* is efficiently disrupted by a muscular grinder in the pharynx of the worm. However, very few intact bacteria may escape from this defense system, and enter in the lumen of the worm intestine (Gupta & Singh 2017). The intestinal lumen of worms is frequently distended during aging, which accompanied by bacterial proliferation (Garigan *et al*. 2002; McGee *et al*. 2011). Blockage of bacterial proliferation by treatment of UV, antibiotics, and heat extends lifespan of worms (Garigan *et al*. 2002; De Arras *et al*. 2014; Hwang *et al*. 2014), implicating that progressive intestinal proliferation of *E. coli* probably contributes to worm aging and death. Thus, age-related dysbiosis in *C. elegans* provides a model to study how host responses to altered gut microbiota to maintain fitness (Ezcurra 2018).

Accumulating evidence has indicated that genetic inactivation of *daf-16* accelerates tissue deterioration and shortens lifespan of wild-type (WT) worms grown on *E. coli* OP50 (Garigan *et al*. 2002; Portal-Celhay & Blaser 2012; Portal-Celhay *et al*. 2012; Li *et al*. 2019). The observation that DAF-16 is activated during bacterial accumulation in older worms (Li *et al*. 2019), prompt us to investigate the role of DAF-16 in dysbiosis in the gut of worms. We found that activation of DAF-16 was required for maintenance of longevity and organismal fitness in worms, at least in part, by up-regulating two lysozyme genes (*lys-7* and *lys-8*), thus limiting bacterial accumulation in the gut of worms during aging. Meanwhile, we identified that indole produced by *E. coli* was involved in the activation of DAF-16 by TRPA-1.

## Results

### Activation of DAF-16 is required for normal lifespan and organismal fitness in worms

To study the role of DAF-16 in age-related dysbiosis, all the experiments started from the young adult stage, which was considered Day 0 (0 days) (Figure 1-figure supplement 1A). Consistent with a recent observation that DAF-16 is activated in worms with age (Li *et al*. 2019), we found that DAF-16::GFP was mainly located in the cytoplasm of the intestine in worms expressing *daf-16p::daf-16::gfp* fed live *E. coli* OP50 on Day 1 (Figure 1A and 1B). The nuclear translocation of DAF-16 in the intestine was increased in worms fed live *E. coli* OP50 on Days 4 and 7, but not in age-matched WT worms fed heat-killed (HK) *E. coli* OP50 (Figure 1A and 1B). To further confirm these results, we tested the expression of two DAF-16 target genes *dod-3* and *hsp-16.2* via the transcriptional reporter strains of *dod-3p::gfp* and *hsp-16.2p::nCherry*. As expected, the expression of either *dod-3p::gfp* or *hsp-16.2p::nCherry* was significantly up-regulated in worms fed live *E. coli* OP50 on Day 4, but not in age-matched worms fed HK *E. coli* OP50 (Figure 1-figure supplement 1B-1D). Likewise, DAF-16 was also retained in the cytoplasm of the intestine in worms fed ampicillin-killed *E. coli* OP50 on Days 4 and 7 (Figure 1-figure supplement 2A). In contrast, starvation induced the nuclear translocation of DAF-16 in the intestine of worms on Day 1 (Figure 1-figure supplement 2B). Thus, either HK or antibiotic-killed *E. coli* OP50 as a food source does not induce a starvation state in worms. Taken together, these results indicate that activation of DAF-16 is mainly attributed to accumulation of live *E. coli*, but not by age itself, in worms.

**Figure 1.**
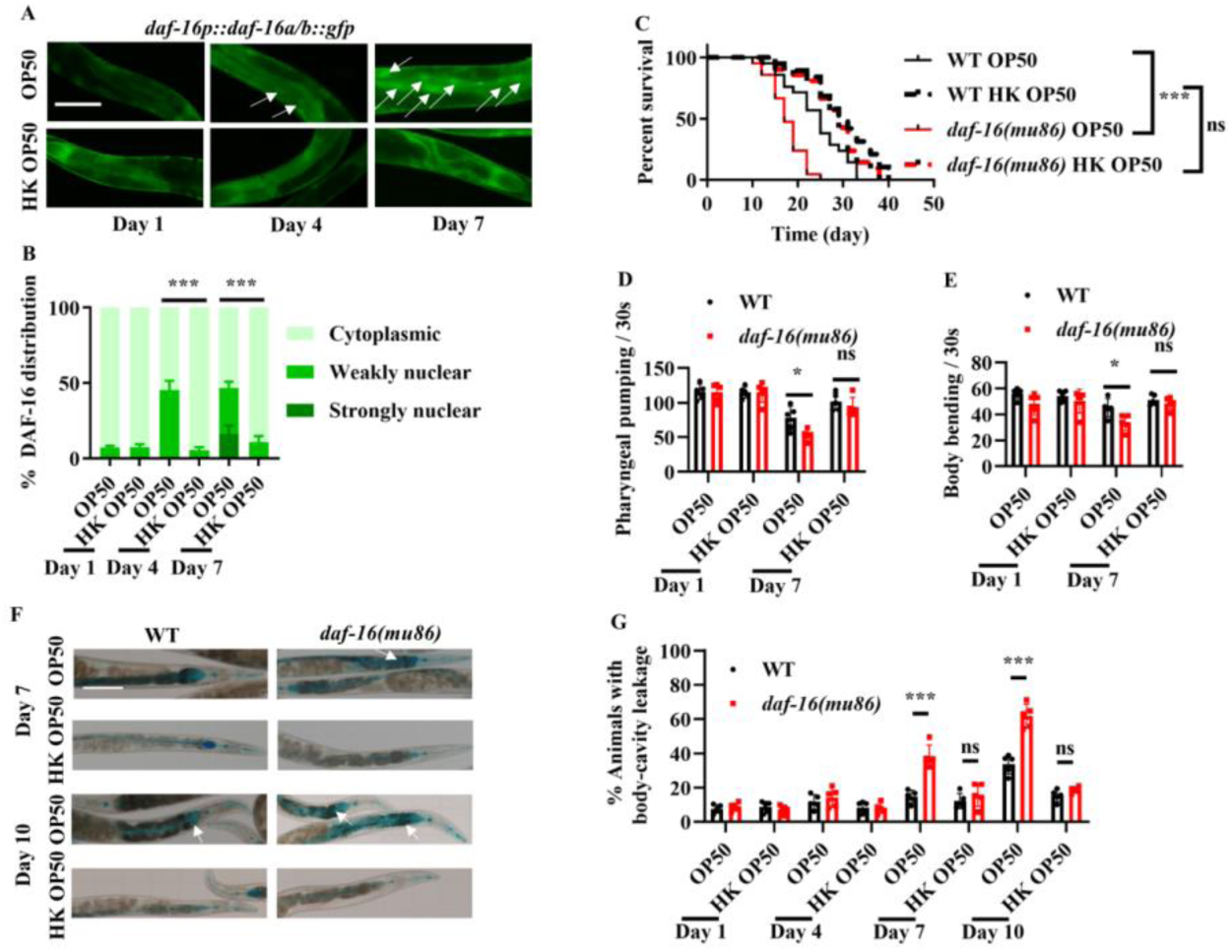
Activation of DAF-16 by bacterial accumulation is required for maintenance of longevity and organismal fitness. **(A)** The nuclear translocation of DAF-16::GFP in the intestine was increased in worms fed live *E. coli* OP50, but not in worms fed heat-killed (HK) *E. coli* OP50. White arrows indicate nuclear localization of DAF-16::GFP. Scale bars: 50 μm. **(B)** Quantification of DAF-16 nuclear localization. These results are means ± SEM of three independent experiments (n > 35 worms per experiment). \*\*\**P* < 0.001. *P*-values were calculated using the using the two-way ANOVA. **(C)** *daf-16(mu86)* mutants grown on live *E. coli* OP50 had a shorter lifespan compared to those grown on HK *E. coli* OP50. \*\*\**P* < 0.001. ns, not significant. *P*-values were calculated using a Log-rank test. **(D and E)** DAF-16 is involved in delaying the appearance of the aging markers, including pharyngeal pumping (D) and body bending (E), in worms fed live *E. coli* OP50, but not in worms fed HK *E. coli* OP50. These results are means ± SEM of five independent experiments (n > 20 worms per experiment). \**P* < 0.05. ns, not significant. (F) Representative images of intestinal permeability stained by food dye FD&C Blue No. 1 in worms. White arrows indicate the body-cavity leakages of worms. Scale bars: 100 μm. **(G)** Quantification of body-cavity leakages was measured in animals fed on live *E. coli* OP50 or heat-killed (HK) *E. coli* OP50 over time. These results are means ± SEM of five independent experiments (n > 20 worms per experiment). \*\*\**P* < 0.001. ns, not significant. *P*-values (**D, E, and G**) were calculated using the unpaired t-test. **Figure 1-source data 1** **Lifespan assays summary and quantification results.**

Consistent with the previous observations (Portal-Celhay *et al*. 2012; Li *et al*. 2019), we found that a mutation in *daf-16(mu86)* shortened lifespan of worms fed live *E. coli* OP50 at 20 °C (Figure 1C). In contrast, the mutation in *daf-16(mu86)* had no impact on lifespan of worms fed either HK (Figure 1C) or ampicillin-killed *E. coli* OP50 (Figure 1-figure supplement 2C). Next, we examined the effects of DAF-16 on phenotypic traits, such as the pharyngeal-pumping rate, body bending, and integrity of intestinal barrier, which are associated with aging in worms. Both the rates of pharyngeal-pumping (Figure 1D) and body bending (Figure 1E) were reduced in *daf-16(mu86)* mutants on Day 7 as compared to those in WT worms fed live *E. coli* OP50. In contrast, the rates of pharyngeal-pumping and body bending were comparable in WT worms and *daf-16(mu86)* mutants grown on HK *E. coli* OP50 on Day 7. Furthermore, we used food dye FD&C Blue No. 1 to evaluate the integrity of intestinal barrier (Ma *et al*. 2020). The body-cavity leakage in *daf-16(mu86)* mutants on Days 7 and 10 were higher than those in age-matched WT worms fed live *E. coli* OP50, but not HK *E. coli* OP50 (Figure 1F and 1G). Taken together, these results demonstrate that activation of DAF-16 by bacterial accumulation is required for maintenance of longevity and organismal fitness.

### Indole produced from *E. coli* activates DAF-16

As DAF-16 is activated by bacterial accumulation, we hypothesized that this activation is probably due to bacterially produced compounds. To test this idea, culture supernatants from *E. coli* OP50 were collected, and freeze-dried. After dissolved in methanol, the crude extract was isolated by high performance liquid chromatograph (HPLC) with automated fraction collector using Agilent ZORBAX SB-C18 liquid chromatography column. A candidate compound was detected by activity-guided isolation, and further identified as indole with mass spectrometry and NMR data (Figure 2A, Figure 2-figure supplement 1A and 1B; Table S1). The observation that indole secreted by *E. coli* OP50 could activate DAF-16 was further confirmed by analyzing commercial HPLC grade indole. Supplementation with indole (50-200 μM) not only significantly induced nuclear translocation of DAF-16 (Figure 2B), but also up-regulated the expression of either *dod-3p::gfp* or *hsp-16.2p::nCherry* in young adult worms after 24 h of treatment (Figure 2-figure supplement 2A and 2B). Next, we found that the levels of indole were 30.9, 71.9, and 105.9 nmol/g dry weight, respectively, in worms fed live *E. coli* OP50 on Days 1, 4, and 7 (Figure 2C). The elevated indole levels in worms were accompanied by an increase in colony-forming units (CFU) of live *E. coli* OP50 in the intestine of worms with age (Figure 2C), suggesting that accumulation of live *E. coli* OP50 in the intestine is probably responsible for increased indole in worms. These data also raised a possibility that exogenous indole produced by *E. coli* OP50 on the NGM plates could increase the levels of indole in worms with age. However, we found that the levels of indole were 28.2, 31.6, and 36.1 nmol/g dry weight, respectively, in worms fed HK *E. coli* OP50 on Days 1, 4, and 7 (Figure 2-figure supplement 3A), indicating that indole was not accumulated in worms fed HK *E. coli* OP50 even for 7 days. Thus, the increase in the levels of indole in worms results from intestinal accumulation of live *E. coli* OP50, rather than exogenous indole produced by *E. coli* OP50 on the NGM plates. The observation that DAF-16 was retained in the cytoplasm of the intestine in worms fed live *E. coli* OP50 on Day 1 (Figure 1A and 1B) also indicated that exogenous indole produced by *E. coli* OP50 on the NGM plates is not enough to activate DAF-16. Supplementation with indole (50-200 μM) significantly increased the indole levels in young adult worms on Day 1 (Figure 2-figure supplement 3B), which could induce nuclear translocation of DAF-16 in worms (Figure 2B).

**Figure 2.**
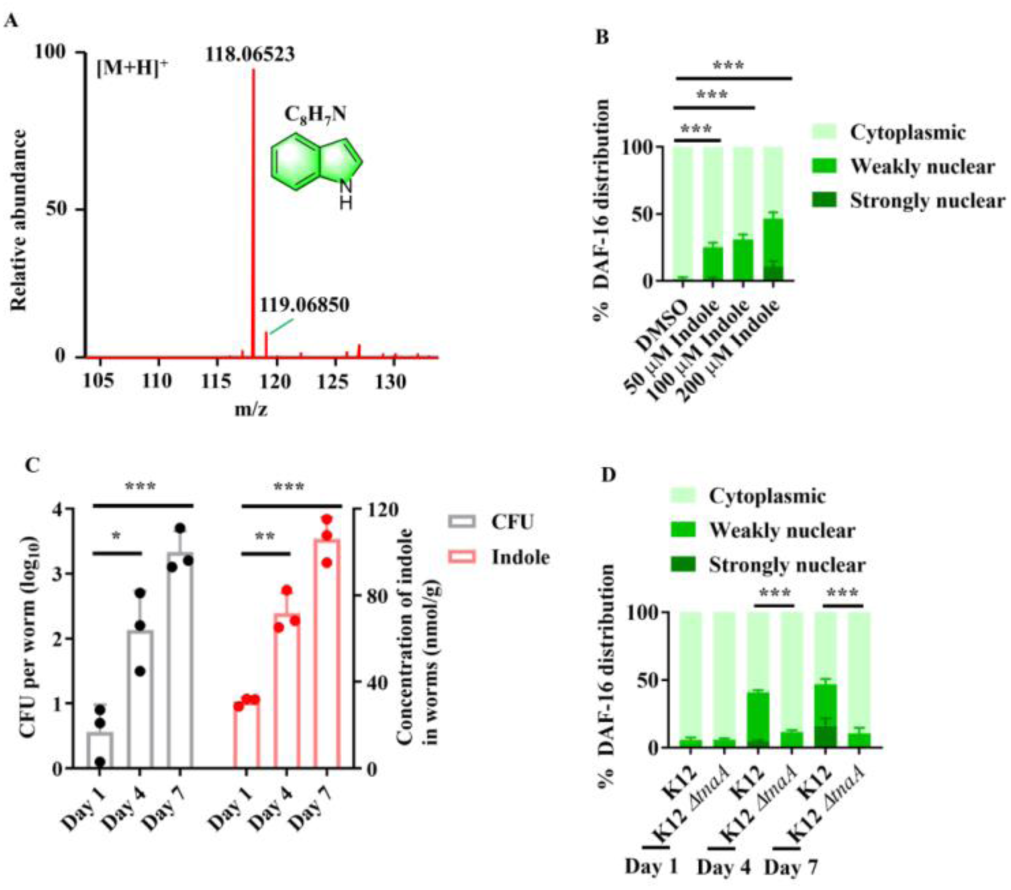
Indole is involved in the nuclear translocation of DAF-16 in worms with age. **(A)** High-resolution mass spectrum of indole. The corresponding 1H and13C NMR spectra were depicted in Figure 2-figure supplement 1 and Table S1. **(B)** Supplementation with indole promoted nuclear translocation of DAF-16::GFP in the intestine of worms. These results are means ± SEM of three independent experiments (n > 35 worms per experiment). \*\*\**P* < 0.001. **(C)** Colony-forming units (CFU) of *E. coli* OP50 were increased in worms over time, which was accompanied by an increase in the levels of indole in worms. These results are means ± SEM of three independent experiments (n > 30 worms per experiment). \**P* < 0.05; \*\**P* < 0.01; \*\*\**P* < 0.001. *P*-values **(C)** were calculated using the unpaired t-test. **(D)** Deletion of *tnaA* significantly suppressed the nuclear translocation of DAF-16::GFP in the intestine of worms fed *E. coli* BW25113. These results are means ± SEM of three independent experiments (n > 35 worms per experiment). \*\*\**P* < 0.001. *P*-values (**B and D**) were calculated using the two-way ANOVA. **Figure 2-source data 1** **Quantification results.**

In bacteria, indole is biosynthesized from tryptophan by tryptophanase (*tnaA*) (Lee *et al*. 2015). To determine the effect of endogenous indole, worms were fed *E. coli* K-12 BW25113 strain (called K-12), and *tnaA*-deficient strain BW25113 *ΔtnaA* (called K-12 *ΔtnaA*), respectively. We found that both *tnaA* mRNA and indole levels were undetectable in the K-12 *ΔtnaA* strain (Figure 2-figure supplement 4A and 4B). Furthermore, disruption of *tnaA* significantly suppressed the nuclear translocation of DAF-16 (Figure 2D). The nuclear translocation of DAF-16::GFP was mainly located in the cytoplasm of the intestine in worms fed live K-12 *ΔtnaA* strains on Day 4. However, supplementation with indole induced the nuclear translocation of DAF-16::GFP in the intestine in these worms (Figure 2-figure supplement 4C). Taken together, our results suggest that indole is involved in the activation of DAF-16 in worms with age.

### Endogenous indole is required for normal lifespan

It has been shown that exogenous indole extends lifespan in *C. elegans* at 16 °C (Sonowal *et al*. 2017). We found that adult worms fed *E. coli* K-12 *ΔtnaA* strains exhibited a shortened lifespan at 20 °C, compared with those fed *E. coli* K-12 strain (Figure 3A). Supplementation with indole in adults not only rescued the shortened lifespan of WT worms fed *E. coli* K-12 *ΔtnaA* strain, but also significantly extended the lifespan of WT worms fed *E. coli* K-12 strain (Figure 3A). In contrast, the lifespan of *daf-16(mu86)* mutants fed *E. coli* K-12 *ΔtnaA* strain was comparable to that of *daf-16(mu86)* mutants fed *E. coli* K-12 strain (Figure 3B). Supplementation with indole (100 μM) did not affect the lifespan of *daf-16(mu86)* mutants fed either *E. coli* K-12 or K-12 *ΔtnaA* strain (Figure 3B). Moreover, the CFU of *E. coli* K-12 *ΔtnaA* strain were significantly higher than those of *E. coli* K12 strain in WT worms on Days 4 and 7 (Figure 3C). By contrast, the CFU of *E. coli* K-12 *ΔtnaA* strain were similar to those of *E. coli* K12 strain in *daf-16(mu86)* mutants on Days 4 and 7 (Figure 3D). Likewise, the accumulation of *E. coli* K-12 *ΔtnaA* strain expressing mCherry were significantly higher than that of *E. coli* K12 strain in WT worms, but not *daf-16(mu86)* mutants, on Days 4 and 7 (Figure 3E-3G). Finally, supplementation with indole (100 μM) inhibited the CFU of *E. coli* K-12 in WT worms, but not *daf-16(mu86)* mutants, on Days 4 and 7 (Figure 3H and 3I). These results suggest that endogenous indole is involved in maintaining normal lifespan in worms.

**Figure 3.**
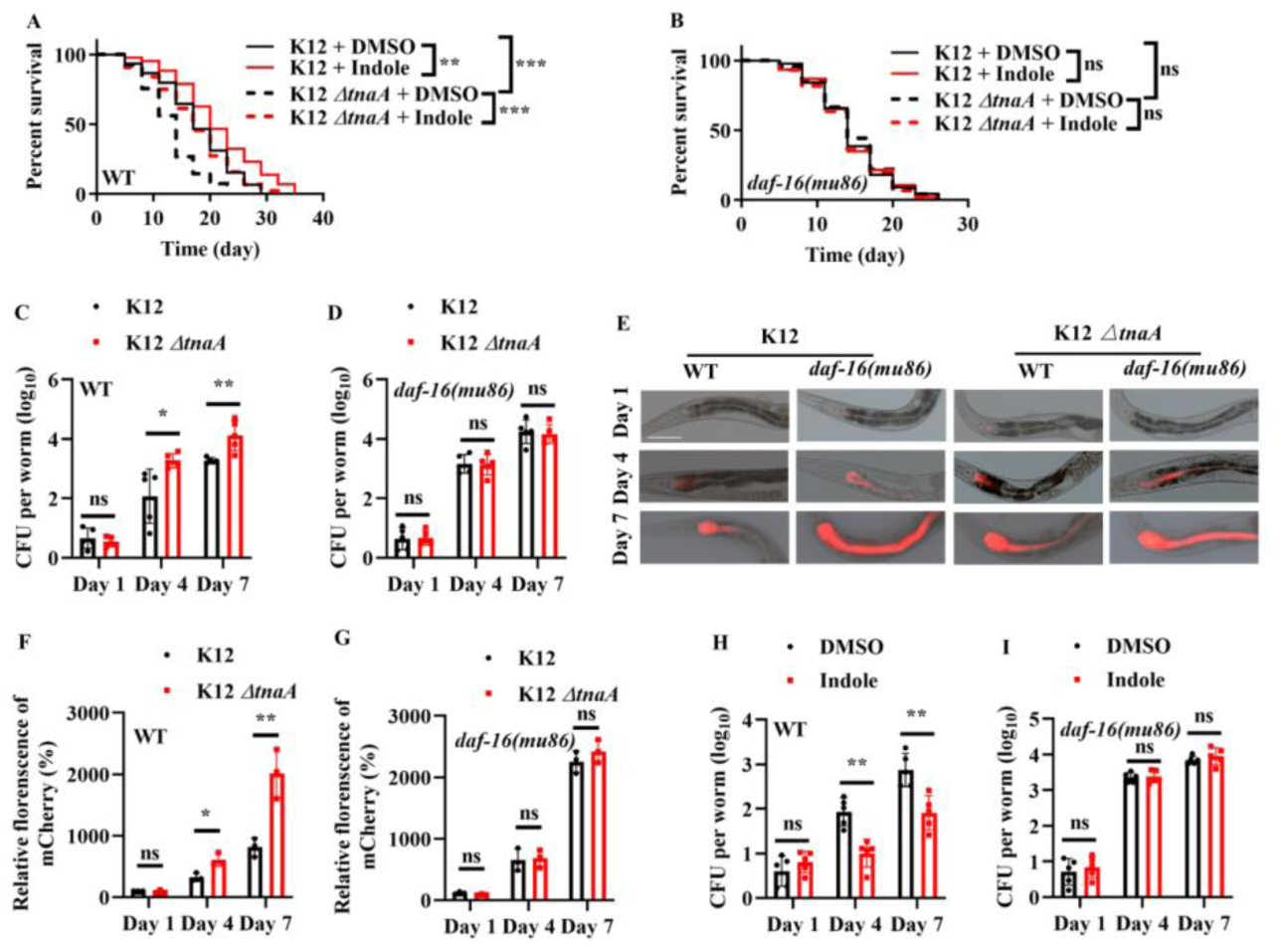
Indole is required for maintenance of normal lifespan via DAF-16 in worms. **(A)** Wild type (WT) worms fed *E. coli* K-12 *ΔtnaA* strains had a shorter lifespan compared to those fed *E. coli* K-12 strain at 20 °C. Supplementation with indole (100 μM) extended lifespan of WT worms fed *E. coli* K-12, and rescued the short lifespan of WT worms fed *E. coli* K-12 *ΔtnaA* strain. \*\**P* < 0.01. \*\*\**P* < 0.001. **(B)** Indole-mediated lifespan extension depended on DAF-16 in worms. ns, not significant. *P*-values (**A and B)** were calculated using log-rank test. **(C and D)** Colony-forming units (CFU) of *E. coli K-12* or K-12 *ΔtnaA* were measured in WT worms (C) or *daf-16(mu86)* mutants (D). These results are means ± SEM of five independent experiments (n > 30 worms per experiment). \**P* < 0.05; \*\**P* < 0.01. ns, not significant. **(E)** Fluorescence images of worms exposed to *E. coli* K-12 or K-12 *ΔtnaA* expressing mCherry. Scale bars: 50 μm. (**F and G**) Quantification of fluorescent intensity of *E. coli* K-12 or K-12 *ΔtnaA* expressing mCherry in WT worms (F) or *daf-16(mu86)* mutants (G). These results are means ± SEM of three independent experiments (n > 35 worms per experiment). \**P* < 0.05. \*\**P* < 0.01. ns, not significant. **(H and I)** CFU of *E. coli* K-12 were measured in WT worms (H) or *daf-16(mu86)* mutants (I) in the presence of exogenous with indole (100 μM). These results are means ± SEM of five independent experiments (n > 30 worms per experiment). \*\**P* < 0.01. ns, not significant. *P*-values (**C, D, and F-I**) were calculated using the unpaired t-test. **Figure 3-source data 1** **Lifespan assays summary and quantification results.**

### TRPA-1 in neurons is required for indole-mediated longevity

How did worms detect bacterially produced indole during aging? Previously, Sonowal et al. have identified that *C. elegans* xenobiotic receptor AHR-1, which encodes an ortholog of the mammalian AHR, mediates indole-promoted lifespan extension in worms at 16 °C (Sonowal *et al*. 2017). However, we found that RNAi knockdown of *ahr-1* did not affect the nuclear translocation of DAF-16 in worms fed *E. coli* K12 strain on Day 7 (Figure 4-figure supplement 1A) or young adult worms treated with indole (100 μM) for 24 h (Figure 4-figure supplement 1B). A recent study has demonstrated that bacteria-derived indole activates the transient receptor potential ankyrin 1 (TRPA1), a cold-sensitive TRP channel, in enteroendocrine cells in zebrafish and mammals (Ye *et al*. 2021). As *C. elegans* TRPA-1 is an ortholog of mammalian TRPA1 (Kindt *et al*. 2007; Venkatachalam & Montell 2007), we tested the role of TRPA-1 in DAF-16 activation by indole. We found that RNAi knockdown of *trpa-1* significantly inhibited the nuclear translocation of DAF-16 in worms fed *E. coli* K12 strain on Days 4 and 7 (Figure 4A) or young adult worms treated with indole (100 μM) for 24 h (Figure 4B). These results suggest that TRPA-1 is involved in indole-mediated DAF-16 activation. Previously, Xiao et al. (Xiao *et al*. 2013) have demonstrated that TRPA-1 activated by low temperatures mediates calcium influx, which in turn stimulates the PKC-2-SGK-1 signaling to promote the transcription activity of DAF-16, leading to lifespan extension in worms. However, we found that RNAi knockdown of *sgk-1* did not influence the nucleocytoplasmic distribution of DAF-16 in the presence of indole (Figure 4-figure supplement 1C). Mutant worms lacking *trpa-1* exhibited a shorter lifespan than did WT worms at 20 °C (Xiao *et al*. 2013). Consistent with this observation, knockdown of *trpa-1* by RNAi significantly shortened the lifespan in worms fed *E. coli* K12 strain (Figure 4C). Supplementation with indole no longer extended the lifespan of worms after knockdown of *trpa-1* by RNAi. Furthermore, the CFU of *E. coli* K-12 strain were significantly increased in worms subjected to *trpa-1* RNAi on Days 4 and 7 (Figure 4D and 4E). Supplementation with indole failed to suppress the increases in CFU in these worms. These results suggest that indole exhibits its function in extending lifespan and inhibiting bacterial accumulation primarily via TRPA-1. It has been shown that podocarpic acid, a TRPA-1 agonist, activates the SEK-1/PMK-1/SKN-1 pathway (Chaudhuri *et al*. 2016), a signaling cascade involved in *C. elegans* defense against pathogenic bacteria (Kim *et al*. 2002). However, we found that supplementation with 0.1 mM indole failed to induce nuclear localization of SKN-1::GFP in the intestine of the transgenic worms expressing *skn-1p::skn-1::gfp* (Figure 4-figure supplement 2A and 2B), suggesting that indole cannot activate SKN-1 in worms.

**Figure 4.**
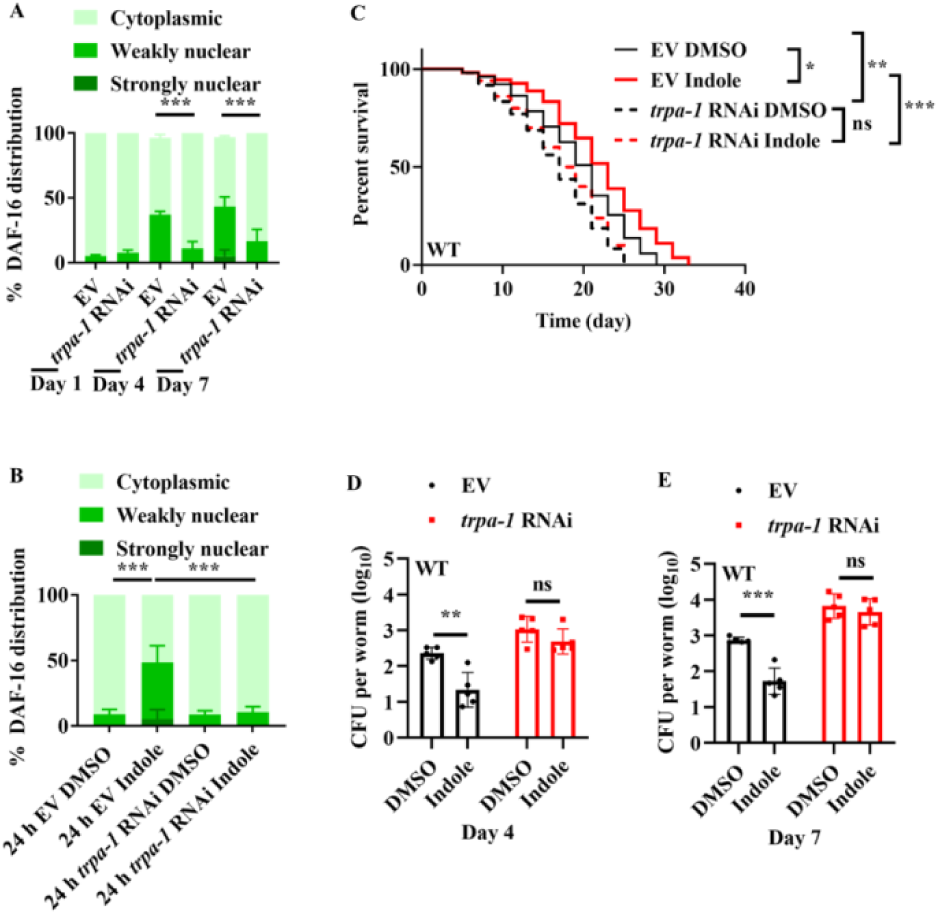
TRPA-1 is involved in indole-mediated DAF-16 activation. **(A and B)** Knockdown of *trpa-1* by RNAi suppressed nuclear translocation of DAF-16::GFP in worms on Days 4 and 7 (A), or in young adult worms treated with indole (100 μM) for 24 h (B). EV, empty vector. These results are means ± SEM of three independent experiments (n > 35 worms per experiment). \*\*\**P* < 0.001. ns, not significant. *P*-values (**A and B**) were calculated using the two-way ANOVA. **(C)** Knockdown of *trpa-1* by RNAi significantly shortened the lifespan in worms treated with indole (100 μM). \**P* < 0.05; \*\**P* < 0.01; \*\*\**P* < 0.001. *P*-values were calculated using log-rank test. **(D and E)** Colony-forming units (CFU) of *E. coli* K12 were significantly increased in worms subjected to *trpa-1* RNAi on Days 4 (D) and 7 (E). Meanwhile, supplementation with indole (100 μM) failed to suppress the increases in CFU in *trpa-1* (RNAi) worms. These results are means ± SEM of five independent experiments (n > 30 worms per experiment). \*\**P* < 0.01; \*\*\**P* < 0.001. ns, not significant. *P*-values (**D** and **E**) were calculated using the unpaired t-test. **Figure 4-source data 1** **Lifespan assays summary and quantification results.**

As overexpression of *trpa-1* in the intestine and neurons was sufficient to extend the lifespan of worms (Xiao *et al*. 2013), we determined tissue-specific activities of TRPA-1 in the regulation of longevity mediated by indole. Consistent with this observation (Xiao *et al*. 2013), we found that both neuronal- and intestinal-specific knockdown of *trpa-1* by RNAi significantly shortened the lifespans in worms (Figure 5A and 5B). However, supplementation with indole (100 μM) only extended the lifespan in worms subjected to intestinal-specific (Figure 5B), but not neuronal-specific, *trpa-1* RNAi (Figure 5A). Likewise, knockdown of *trpa-1* in either neurons (Figure 5C and 5D) or the intestine (Figure 5E and 5F) increased the CFU of *E. coli* K-12 in worms on Days 4 and 7. However, supplementation with indole failed to inhibit the increase in the CFU of *E. coli* K-12 in worms subjected to neuronal-specific *trpa-1* RNAi (Figure 5C and 5D). By contrast, supplementation with indole significantly suppressed the CFU of *E. coli* K-12 in these worms subjected to intestinal-specific *trpa-1* RNAi (Figure 5E and 5F). These results suggest that TRPA-1 in neurons is involved in indole-mediated longevity.

**Figure 5.**
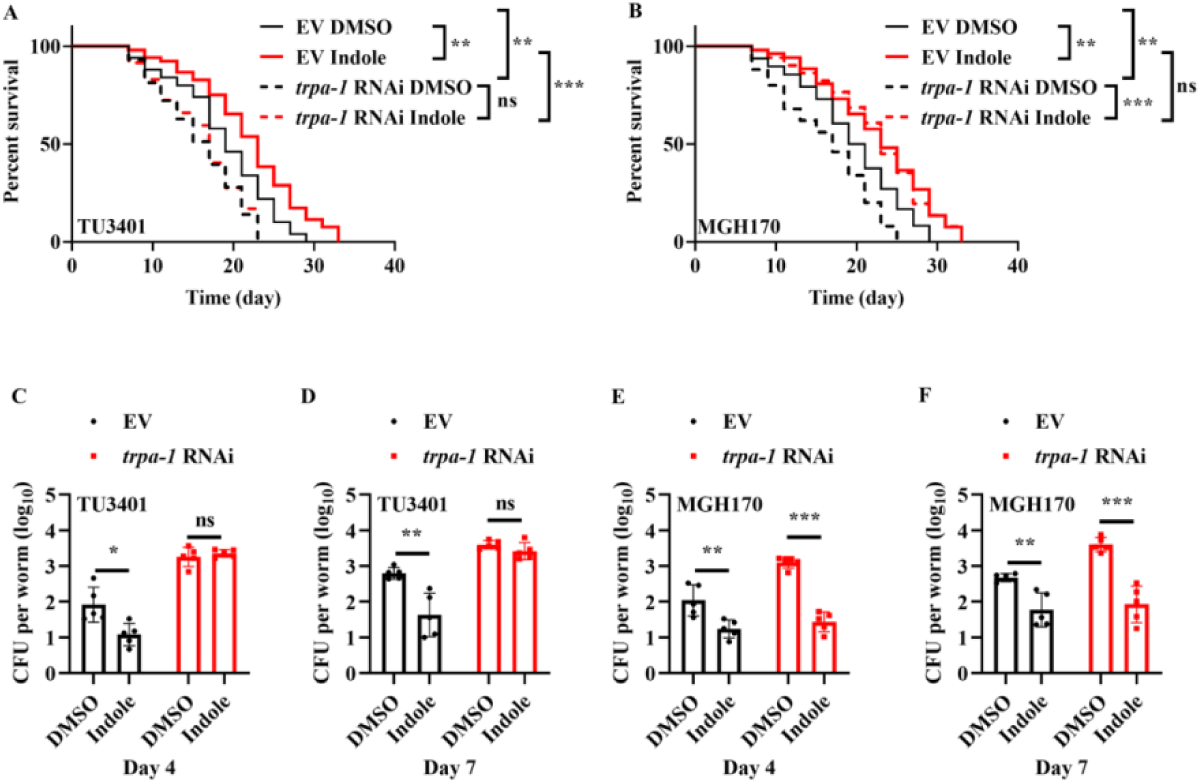
TRPA-1 in neurons is involved in indole-mediated longevity. **(A and B)** Either neuronal - (A) or intestinal -(B) specific *trpa-1* RNAi shortened lifespan in worms. However, supplementation with indole (100 μM) significantly extended lifespan in worms after knockdown of *trpa-1* by RNAi in the intestine, but not in neurons. EV, empty vector. \*\**P* < 0.01; \*\*\**P* < 0.001. ns, not significant. *P*-values (**A** and **B**) were calculated using log-rank test. **(C and D)** Supplementation with indole (100 μM) no longer inhibited colony-forming units (CFU) of *E. coli* K12 in worms on Days 4 (C) and 7 (D) after knockdown of *trpa-1* by RNAi in neurons. These results are means ± SEM of five independent experiments (n > 30 worms per experiment). \**P* < 0.05; \*\**P* < 0.01. ns, not significant. **(E and F)** Supplementation with indole (100 μM) significantly suppressed the CFU of *E. coli* K-12 in worms subjected to intestinal-specific *trpa-1* RNAi on Days 4 (E) and 7 (F). These results are means ± SEM of five independent experiments (n > 30 worms per experiment). \*\**P* < 0.01; \*\*\**P* < 0.001. ns, not significant. *P*-values (**C-F**) were calculated using the unpaired t-test. **Figure 5-source data 1** **Lifespan assays summary and quantification results.**

### LYS-7 and LYS-8 functions as downstream molecules of DAF-16 to maintain normal lifespan in worms

*C. elegans* possesses a variety of putative antimicrobial effector proteins, such as lysozymes, defensin-like peptides, neuropeptide-like proteins, and caenacins (Dierking *et al*. 2016). Of these antimicrobial proteins, *C. elegans* lysozymes are involved in host defense against various pathogens (Mallo *et al*. 2002; O’Rourke *et al*. 2006; Irazoqui *et al*. 2010a; Boehnisch *et al*. 2011; Visvikis *et al*. 2014). A previous study has demonstrated that expressions of lysozyme genes, such as *lys-2, lys-7,* and *lys-8*, are markedly up-regulated in 4-day-old worms, which is dependent on DAF-16 (Li *et al*. 2019). We thus determined the role of these lysozyme genes in lifespan of worms. We found that a single mutation in *lys-7(ok1384)* slightly but significantly reduced the lifespan in worms fed live *E. coli* OP50 (Figure 6-figure supplement 1A), but not HK *E. coli* OP50 (Figure 6-figure supplement 1B), which was consistent with a previous observation (Portal-Celhay *et al*. 2012). In contrast, whereas either a single mutation in *lys-8(ok3504)* or RNAi knockdown of *lys-2* did not affect the lifespan in worms grown on live *E. coli* OP50, or HK *E. coli* OP50 (Figure 6-figure supplement 1A and 1B). However, *lys-7(ok1384); lys-8(ok3504)* double mutants exhibited a shortened lifespan in worms fed live *E. coli* OP50 (Figure 6A). In contrast, the lifespan of *lys-7(ok1384); lys-8(ok3504)* double mutants was comparable to that of WT worms fed HK *E. coli* OP50 (Figure 6B). Furthermore, knockdown of *lys-2* by RNAi did not affect the lifespan of *lys-7(ok1384); lys-8(ok3504)* double mutants grown on live *E. coli* OP50, or HK *E. coli* OP50 (Figure 6-figure supplement 1C and 1D). Both the rates of pharyngeal-pumping (Figure 6C) and body bending (Figure 6D) were reduced in 8-day-old *lys-7(ok1384); lys-8(ok3504)* double mutants fed live *E. coli* OP50. Moreover, the body-cavity leakage was increased in the double mutants fed live *E. coli* OP50 on Day 10 (Figure 6E). However, these age-associated markers in *lys-7(ok1384); lys-8(ok3504)* double mutants were comparable to those in in age-matched WT worms fed HK *E. coli* OP50. In addition, we found that double mutations in *lys-7(ok1384); lys-8(ok3504)* increased the CFU of *E. coli* OP50 (Figure 6F) as well as accumulation of *E. coli* OP50 expressing RFP in worms on Day 7 (Figure 6G). Finally, we found that supplementation with indole no longer extended lifespan and failed to suppress the increase the CFU of *E. coli* K-12 in *lys-7(ok1384); lys-8(ok3504)* double mutants (Figure 6-figure supplement 2A and 2B).

**Figure 6.**
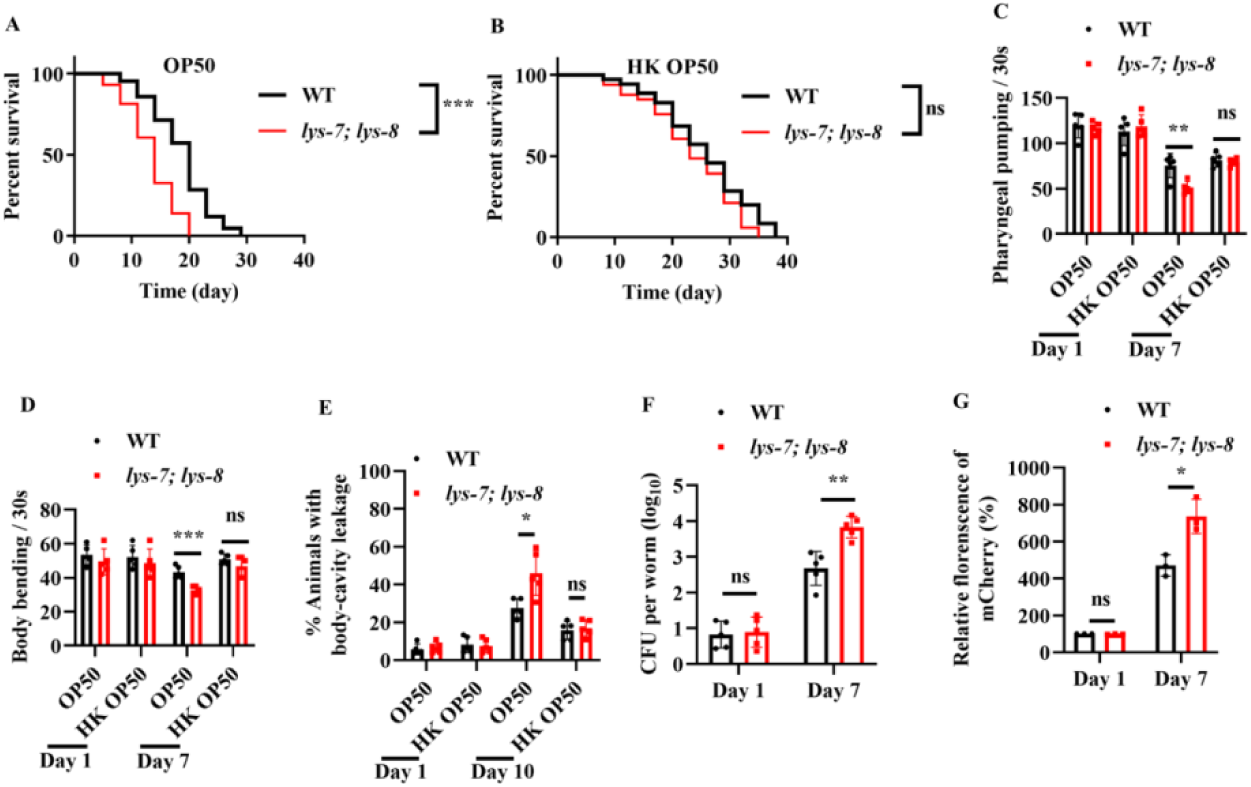
LYS-7 and LYS-8 are required for maintenance of normal lifespan in worms. **(A and B)** The lifespans of worms fed either live *E. coli* OP50 (A) or fed heat-killed (HK) *E. coli* OP50 (B). The *lys-7(ok1384); lys-8(ok3504)* double mutants exhibited a shorter lifespan compared to wild-type (WT) worms fed live *E. coli* OP50 (A). By contrast, the lifespan in the *lys-7(ok1384); lys-8(ok3504)* double mutants was comparable of that in WT worms fed HK *E. coli* OP50 (B). \*\*\**P* < 0.001. ns, not significant. *P*-values (**A** and **B**) were calculated using log-rank test. **(C-E)** *lys-7* and *lys-8* were involved in delaying the appearance of the aging markers, including pharyngeal pumping (C), body bending (D), and body-cavity leakage (E) in worms fed live *E. coli* OP50. These results are means ± SEM of five independent experiments (n > 20 worms per experiment). \**P* < 0.05; \*\**P* < 0.01; \*\*\**P* < 0.001. **(F)** Colony-forming units (CFU) of *E. coli* OP50 were significantly increased in *lys-7(ok1384)*;*lys-8(ok3504)* double mutants on Day 7. These results are means ± SEM of five independent experiments (n > 20 worms per experiment). \*\**P* < 0.01. ns, not significant. **(G)** Quantification of fluorescent intensity of *E. coli* OP50 expressing mCherry in *lys-7(ok1384);lys-8(ok3504)* double mutants. These results are means ± SEM of three independent experiments (n > 35 worms per experiment). \**P* < 0.05. ns, not significant. *P*-values (**C-G**) were calculated using the unpaired t-test. **Figure 6-source data 1** **Lifespan assays summary and quantification results**.

Using the transgenic worms expressing either *lys-7p::gfp* or *lys-8p::gfp*, we found that the expressions of *lys-7p::gfp* and *lys-8p::gfp* were significantly up-regulated in worms fed live *E. coli* OP50 on Days 4 and 7 (Figure 7-figure supplement 1A-1D), but not in age-matched worms fed HK *E. coli* OP50 (Figure 7-figure supplement 1A-1D). RNAi knockdown of *daf-16* also significantly suppressed the expressions of *lys-7p::gfp* and *lys-8p::gfp* in these worms fed live *E. coli* OP50 (Figure 7A and 7B). Similar results were obtained by measuring the mRNA levels of *lys-7* and *lys-8* in *daf-16(mu86)* mutants using quantitive real-time PCR (qPCR) (Figure S11E and S11F). Furthermore, we found that expression of either *lys-7p::gfp* (Figure 7C) or *lys-8p::gfp* (Figure 7D) was reduced in worms fed *E. coli* K-12 *ΔtnaA* strain on Days 4 and 7, compared to those in age-matched worms fed *E. coli* K-12 strain. Likewise, indole (100 μM) remarkably increased the expression of either *lys-7p::gfp* or *lys-8::gfp* in young adult worms after 24 h treatment (Figure 7E and 7F). Finally, we found that RNAi knockdown of *trpa-1* significantly inhibited the expression of either *lys-7p::gfp* or *lys-8p::gfp* in 4-day-old worms fed *E. coli* K12 strain (Figure 7A and 7B). Meanwhile, we found that the mRNA levels of *lys-7* and *lys-8* were significantly down-regulated in worms subjected to neuronal-specific (Figure 7-figure supplement 2A), but not intestinal-specific, knockdown of *trpa-1* by RNAi on Day 4 (Figure 7-figure supplement 2B). However, supplementation with indole only up-regulated the expression of *lys-7* and *lys-8* in worms subjected to intestinal-specific (Figure 7-figure supplement 2C), but not neuronal-specific, RNAi of *trpa-1* (Figure 7-figure supplement 2D). These results suggest that LYS-7 and LYS-8 function as downstream molecules of DAF-16 to maintain normal lifespan by inhibiting bacterial proliferation in worms.

**Figure 7.**
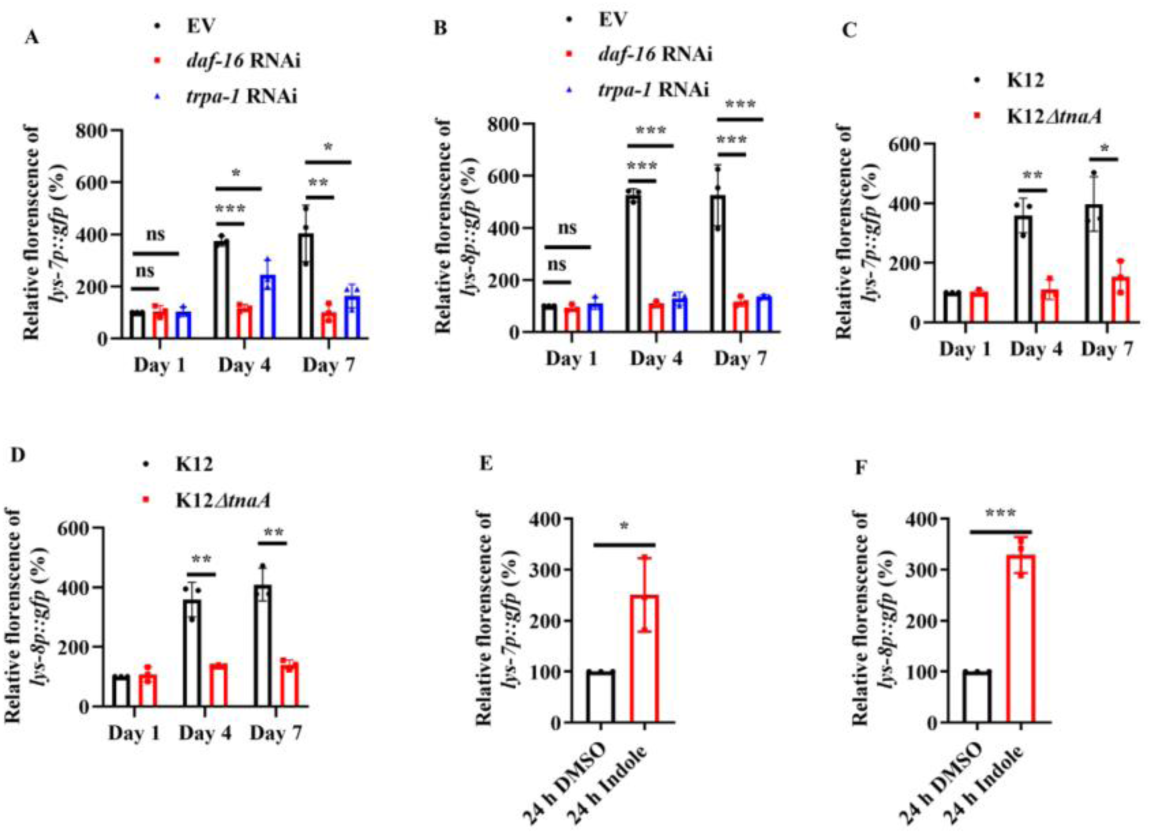
The expressions of *lys-7* and *lys-8* were up-regulated by the indole/TRPA-1/DAF-16 signaling. **(A and B)** The expression of either *lys-7p::gfp* (A) or *lys-8p::gfp* (B) was significantly suppressed after knockdown of *daf-16* or *trpa-1* by RNAi in worms fed live *E. coli* OP50 on Days 4 and 7. EV, empty vector. **(C and D)** The expression of either *lys-7p::gfp* (C) or *lys-8p::gfp* (D) was reduced in worms fed *E. coli* K-12 *ΔtnaA* strain on Days 4 and 7, compared with that in age-matched worms fed *E. coli* K-12. **(E and F)** Indole (100 μM) remarkably increased the expression of either *lys-7p::gfp* (E) or *lys-8::gfp* (F) in young adult worms after 24 h of treatment. These results are means ± SEM of three independent experiments (n > 35 worms per experiment). \**P* < 0.05. \*\**P* < 0.01. \*\*\**P* < 0.001. ns, not significant. *P*-values (**A-H**) were calculated using the unpaired t-test. **Figure 7-source data 1** **Quantification results.**

## Discussion

Using *C. elegans* and its dietary bacterium as a host-microbe model, we provide a striking example of how a host responds to microbial dysbiosis in the gut. As worms age, *E. coli* proliferates in the lumen, thereby producing and secreting more indole. When its concentration crosses a threshold value, the bacterially produced compound is perceived by TRPA-1 in worms. The TRPA-1 signaling triggers DAF-16 nuclear translocation, leading to up-regulation of lysozyme genes. These antimicrobial peptides help worms to maintain normal lifespan by limiting the bacterial proliferation.

A complication in understanding the impact of bacterial diets on the traits of the worm is the fact that initially bacteria are a source of food, but later become pathogenic (Garigan *et al*. 2002; Tan & Shapira 2011). Thus, the accumulation of *E. coli* in the gut is harmful for organismal fitness of worms during the course of life. As a well-known regulator of longevity and innate immunity (Garsin *et al*. 2003; Zou *et al*. 2013), DAF-16 is activated in aging *C. elegans* (Li *et al*. 2019). This transcription factor is involved in both maintaining normal lifespan and limiting proliferation of *E. coli* in worms (Garigan *et al*. 2002; Portal-Celhay & Blaser 2012; Portal-Celhay *et al*. 2012). In this study, our data demonstrate that activation of DAF-16 requires contact with live bacterial cells in the gut of worms as dead *E. coli* fails to activate DAF-16. Thus, accumulation of *E. coli* during aging, but not aging itself, results in the activation of DAF-16. Furthermore, DAF-16 mutation does not influence the lifespan in worms fed dead *E. coli*. Taken together, these findings clearly demonstrate that DAF-16 acts to maintain homeostasis by inhibiting bacterial proliferation in worms with age.

Indole is produced from tryptophan by tryptophanase in a large number of bacterial species. As a well-known signaling molecule, indole is involved in regulation of a variety of physiological processes in bacteria, such as cell division, biofilm formation, virulence, spore formation, and antibiotic resistance (Lee *et al*. 2015; Zarkan *et al*. 2020). Although animals cannot synthesize indole, they can sense and modify this metabolite (Lee *et al*. 2015). Indole and its derivatives can influence insect behaviors and human diseases, such as intestinal inflammation and diabetes (Lee *et al*. 2015; Agus *et al*. 2018). Thus, indole may function as an interspecies and interkingdom signaling molecule to influence the microbe-host interaction (Bansal *et al*. 2010; Oh *et al*. 2012; Ye *et al*. 2021). For instance, enteric delivery of indole (1 mM) increases intestinal motility by inducing 5-hydroxytryptamine secretion in zebrafish larvae (Ye *et al*. 2021). Supplementation with 0.2 mM indole enhances the resistance of *C. elegans* to infection with *C. albicans* by reducing fungal colonization in the intestine (Oh *et al*. 2012). In addition, treatment of HCT-8 intestinal epithelial cells with 1 mM indole inhibits TNF-α-mediated activation of NF-κB and up-regulation of the inflammatory factor IL-8, thus improving intestinal epithelial barrier function (Bansal *et al*. 2010). Our data show that indole limits the bacterial proliferation in the gut of worms by driving intestinal defense gene expression via the transcription factor DAF-16. These findings suggest that the bacteria-derived metabolite may serve as a pathogen-associated molecular pattern that is recognized by metazoans. Although it has been shown that indole at a higher concentration (5 mM) is capable of inhibiting cell division in *E. coli* K12 strain (Chimerel *et al*. 2012), exogenous indole has little effect on the growth of *E. coli* K12 strain up to 3 mM (Chant & Summers 2007). In general, extracellular indole concentrations detected in stationary phase LB cultures are typically 0.5-1 mM depending on the specific *E. coli* strain (Zarkan *et al*. 2020). Our data show that indole is sufficient to inhibit accumulation of *E. coli* BW25113 with the concentration range from 0.05-0.2 mM. Thus, these results suggest that direct causal involvement of cell cycle arrest in *E. coli* by indole is unlikely.

Our data demonstrate that the cold-sensitive TRP channel TRPA-1 in neurons is involved in indole-mediated nuclear translocation of DAF-16 in the intestine, which is required for lifespan extension in *C. elegans*. *trpa-1* is widely expressed in a variety of sensory neurons in worms (Kindt *et al*. 2007). One possibility is that TRPA-1 in neurons may release a neuropeptide, which in turn triggers a signaling pathway to extend lifespan of worms via activating DAF-16 in a non-cell autonomous manner. Low temperature also activates TRPA-1, which in turn acts to promote longevity via the PKC-2-SKG-1-DAF-16 pathway (Xiao *et al*. 2013). It should be noted that unlike indole, activation of TPRA-1 by low temperature promotes the transcription activity, but not the nuclear translocation, of DAF-16 (Xiao *et al*. 2013). Furthermore, our data show that knockdown of *sgk-1* dose not influence the nuclear translocation of DAF-16 induced by indole. Finally, indole extends lifespan via TRPA-1 only in nervous system, whereas low temperature extends lifespan via TRPA-1 both in the intestine and neurons. Thus, indole-mediated lifespan extension is essentially different from low temperature-mediated lifespan extension, although TRPA-1 and DAF-16 are involved in both processes. Although podocarpic acid, the agonist of TRPA-1, can activate the TRPA-1-SKN-1 pathway, SKN-1 is unlikely to be involved in indole-mediated biological effects. We observe that indole fails to activate SKN-1 in worms. More importantly, podocarpic acid does not affect the lifespan in worms (Chaudhuri *et al*. 2016). These data implicate that TRPA-1 activation via indole and via podocarpic acid happens through distinct mechanisms.

In this study, our data demonstrate that DAF-16 inhibits bacterial proliferation by up-regulating expression of two lysozyme genes, *lys-7* and *lys-8*. As a group of digestive enzymes with antimicrobial properties, lysozymes play an important role in the innate immunity in both vertebrate and invertebrate animals (O’Rourke *et al*. 2006). As a target gene of DAF-16, *lys-7* has been proven to play an important role in resistance against a variety of pathogens, such as *Microbacterium nematophilum* (O’Rourke *et al*. 2006), *Pseudomonas aeruginosa* (Nandakumar & Tan 2008), the pathogenic *E. coli* LF82 (Simonsen *et al*. 2011), *Bacillus thuringiensis* (Boehnisch *et al*. 2011), and *Cryptococcus neoformans* (Marsh *et al*. 2011). A previous study has demonstrated that a mutation in *lys-7* significantly reduces lifespan, but does not influence the accumulation of *E. coli* OP50 in the intestine of 2-days-old worms (Portal-Celhay *et al*. 2012). In the current study, the *lys-7* mutants exhibit reduced lifespan and elevated bacterial loads in the intestine of worms on Days 4 and 7. This discrepancy may be due to the different worms at different ages. Actually, our data also show that the *lys-7;lys-8* double mutants exhibit a comparable accumulation of *E. coli* OP50 in worms on Day 1. Although *lys-8* mutation dose not influence either lifespan or bacterial loads, it enhances the effect of *lys-7*. These results suggest that *lys-8* acts in synergy with *lys-7* to limit bacterial accumulation in the gut of worms.

It has been well established that bacterial dysbiosis is significantly associated with inflammatory bowel diseases (IBD) (Manichanh *et al*. 2012; Sommer & Backhed 2013). Interestingly, reduced levels of indole and its derivative indole-3-propionic acid (IPA) are observed in serum of mice with dextran sulfate sodium-induced colitis and patients with IBD (Alexeev *et al*. 2018). Oral administration of IPA significantly ameliorates disease and promotes intestinal homeostasis by up-regulating colonic epithelial IL-10R1 in the chemically induced murine colitis model. Thus, characterization of the role for indole and its derivatives in host–microbiota interactions within the mucosa may provide new therapeutic avenues for inflammatory intestinal diseases.

## Materials and Methods

### Nematode strains

daf-16(mu86), lys-7(ok1384), lys-8(ok3504), LD1[skn-1b/c::gfp + rol-6(su1006)], the nematode strain for neuronal-specific RNAi, TU3401 (sid-1(pk3321); uIs69 [pCFJ90 (myo-2p:: mCherry) + unc-119p::sid-1]), TJ356 (zIs356 [daf-16p::daf-16a/b::GFP + rol-6 (su1006)]), SAL105 (denEx2 [lys-7::GFP + pha-1(+)]) were kindly provided by the Caenorhabditis Genetics Center (CGC; http://www.cbs.umn.edu/CGC), funded by NIH Office of Research Infrastructure Programs (P40 OD010440). The nematode strain for intestinal-specific RNAi, MGH170 (sid-1(qt9); Is[vha-6pr::sid-1]; Is[sur-5pr::GFPNLS]), was kindly provided by Dr. Gary Ruvkun (Massachusetts General Hospital, Harvard Medical School). The strain MQD1586 (476[hsp-16.2p::nCherry; dod-3p::gfp; mtl-1::bfp, unc-119(+)]) was kindly provided by Dr. Mengqiu Dong (Beijing Institute of Life Sciences). Mutants were backcrossed three times into the N2 strain used in the laboratory. All strains were maintained on nematode growth media (NGM) and fed with E. coli OP50 at 20 °C.

### Study Design for age-related dysbiosis study

Synchronized L1 larvae were grown on NGM agar plates seeded with *E. coli* OP50 at 20 °C until they reached the young adult stage. All the experiments started from the young adult stage, which was considered Day 0 (0 days). From Day 1 to Day 10, the worms were transferred to new NGM plates containing *E. coli* strains at 20 °C daily for further experiments. BW21153 (*E. coli* K-12 wild-type) and *E. coli* K-12 *ΔtnaA* strain were obtained from the Keio collection (Baba *et al*. 2006).

### RNA interference

RNAi bacterial strains containing targeting genes were obtained from the Ahringer RNAi library (Kamath & Ahringer 2003). All clones used in this study were verified by sequencing. Briefly, *E. coli* strain HT115 (DE3) expressing dsRNA was grown in LB (Luria-Bertani) containing 100 μg/ml ampicillin at 37 °C for overnight, and then spread onto NGM plates containing 100 μg/ml ampicillin and 5 mM isopropyl 1-thio-β-D-galactopyranoside (IPTG). The RNAi-expressing bacteria were then grown at 25 °C overnight. Synchronized L1 larvae were placed on the plates at 20 °C until they reached maturity. Young adult worms were used for further experiments.

### Construction of transgenic strains

The vector expressing *lys-8p::gfp* was generated by subcloning a 2011bp promoter fragment of *lys-8* into an expression vector (pPD95.75). The vector was injected into the syncytial gonads of WT worms with 50 ng/ml pRF4 as a transformation marker (Mello & Fire 1995). The transgenic worms carrying were confirmed before assay.

### DAF-16 nuclear localization assay

For the effect of aging on DAF-16::GFP localization, worms expressing *daf-16p::daf-16::gfp* were cultured on standard NGM plates at 20 °C for 1, 4, and 7 days, respectively. For indole treatment, young adults were transferred to NGM plates containing 50-200 μM indole (Macklin, Shanghai, China) dissolved in DMSO for 24 h at 20 °C. NGM plates with equal amount of ethanol served as the control. After taken from incubation, worms were immediately mounted in M9 onto microscope slides. The slides were viewed using a Zeiss Axioskop 2 plus fluorescence microscope (Carl Zeiss, Jena, Germany) with a digit camera. The status of DAF-16 localization was categorized as cytosolic localization, nuclear localization when localization is observed throughout the entire body, or intermediate localization when nuclear localization is visible, but not completely throughout the body (Oh *et al*. 2005). At least 35 nematodes were counted in each experiment.

### Fluorescence microscopic analysis

For imaging fluorescence, worms expressing *hsp-16.2p::nCherry*, *dod-3p::gfp*, *lys-7p::gfp*, and *lys-8p::gfp* were mounted in M9 onto microscope slides. The slides were imaged using a Zeiss Axioskop 2 Plus fluorescence microscope. The intensities of nCherry and GFP were analyzed using the ImageJ software (NIH). Three plates of at least 35 animals per plate were tested per assay, and all experiments were performed three times independently.

### Lifespan analysis

Synchronized L1 larvae were grown on NGM agar plates seeded with *E. coli* OP50 at 20 °C until they reached the young adult stage. All the lifespan assays started from the young adult stage at 20 °C. The first day of adulthood was recorded as Day 1. From Day 1 to Day 10, the worms were transferred to new NGM plates containing *E. coli* strains at 20 °C daily. After that, worms were transferred every third day. The number of worms was counted every day. Worms that did not move when gently prodded and displayed no pharyngeal pumping were marked as dead.

### Age-related phenotypic marker assays

The following two age-related phenotypes were scored in 1- and 7-day-old worms (Chen *et al*. 2019). Pharyngeal pumping was measured by counting the number of contractions in the terminal bulb of pharynx in 30 s intervals. Body bending was measured by counting the number of body bends in 30 s intervals. At least 20 animals were determined per assay in 5 independent experiments.

### Intestinal barrier function assay in worms

Intestinal barrier function was determined according to the method described previously (Ma *et al*. 2020). Briefly, synchronized young adult animals were cultured on standard NGM plates at 20 °C for 1, 4, 7, and 10 days. After removed from the NGM plates, these animals were suspended in M9 liquid medium containing *E*. *coli* OP50 (OD = 0.5-0.6), 5 % food dye FD&C Blue No. 1 (Bis[4-(N-ethyl-N-3-sulfophenylmethyl) aminophenyl]-2-sulfophenylmethylium disodium salt) (AccuStandard, New Haven, CT), and incubated for 6 h. After collected and washed with M9 buffer four times, the worms were mounted in M9 onto microscope slides. The slides were viewed using a Zeiss Axioskop 2 Plus fluorescence microscope (Carl Zeiss, Jena, Germany) to measure the leakage of the dyes in the body cavity of animals. The rate of body-cavity leakage was calculated as a percentage by dividing the number of animals with dye leakage by the number of total animals. For each time point, five independent experiments were carried out. In each experiment, at least 20 of worms were calculated.

### Detection of bacterial accumulation in worms

For detection of *E. coli* accumulation, worms were grown on NGM plates with *E. coli* expressing mCherry (The plasmids of PMF440 purchased from addgene) for 1, 4, 7 days at 20 °C. Then animals were collected and soaked in M9 buffer containing 25 mM levamisole hydrochloride (Sangon Biotech Co.), 50 μg/ml kanamycin (Sangon Biotech Co.), and 100 μg/ml ampicillin (Sangon Biotech Co.) for 30 minutes at room temperature. Then worms were washed three times with M9 buffer. Some of animals were mounted in M9 onto microscope slides. The slides were viewed using a Zeiss Axioskop 2 plus fluorescence microscope (Carl Zeiss, Jena, Germany) with a digital camera. At least 35 worms were examined per assay in three independent experiments. Meanwhile, at least 30 of nematodes were transferred into 50 μl PBS plus 0.1 % Triton and ground. The lysates were diluted by 10-fold serial dilutions in sterilized water and spread over LB agar plates with 100 μg/ml ampicillin. After incubation overnight at 37 °C, the *E. coli* CFU was counted. For each group, 3-5 independent experiments were carried out.

### Isolation and identification of the active compound

Ten liters of the *E. coli* OP50 culture supernatants were collected by centrifugation, and freeze-dried in a VirTis freeze dryer. The powders were then dissolved with 10 ml of methanol. The crude extract was loaded on to an Ultimate 3000 HPLC (Thermofisher, Waltham, MA) coupled with automated fraction collector in batches through a continuous gradient on an Agilent ZORBAX SB-C18 column (Agilent, 5 μm, 4.6 × 250 mm) at a column temperature of 40 °C to yield 45 fractions based on retention time and one fraction was collected per minute. The total flow rate was 1 ml/min; mobile phase A was 0.1 % formic acid in water; and mobile phase B was 0.1 % formic acid in acetonitrile. The HPLC conditions were manually optimized on the basis of separation patterns with the following gradient: 0-2 min, 10 % B; 10 min, 25 % B; 30 min, 35 % B; 35 min, 90 % B; 36 min, 95 % B; 40 min, 90 % B; 40.1 min, 10 % B; and 45 min, 10 % B. UV spectra were recorded at 204-400 nm. The injection volume for the extracts was 50 μl. These fractions were tested for detection of nuclear translocation of DAF-16::GFP in worms. We found that the 26th fraction, which was collected at 26-27 min, could induce nuclear translocation of DAF-16 in worms. The active fraction was further then purified by Sephadex LH-20 with methanol and a single candidate active compound was obtained. The purified compound was structurally elucidated with NMR and MS data. NMR experiments were carried out on a Bruker DRX-500 spectrometer (Bruker Corp., Madison, WI) with solvent as internal standard. High-resolution ESI-MS data was performed on Q Exactive Focus UPLC-MS (Thermofisher) with a PDA detector and an Obitrap mass detector using positive mode electrospray ionization.

### Quantitative analysis of indole in worms

For quantitation of indole in *C. elegans*, worms were collected, washed 3-5 times with M9 buffer, and lyophilized for 4-6 h using a VirTis freeze dryer. Dried pellets were weighed, and transferred to a 1.5 ml centrifuge tube. After grinded for 10 minutes in a tissue grinder, the samples were dissolved in 300 µl solvent (methanol: water 20: 80 % (v/v)). The mixtures were then grinded for another 10 minutes, and centrifuged at 12000×g for 5 min. The supernatants were collected, and filtered through 0.22 µm membranes for further analysis using LC-MS. LC-MS analyses were performed on a Q Exactive Focus UPLC-MS (Thermofisher) with an atmospheric pressure chemical ionization (APCI) source and operated with positive mode and coupled with Atlabtis dC18 column (Waters, 3µm, 2.1×150 mm). Five microliters of samples were injected to the LC-MS system for analysis. The total flow rate was 0.3 ml/min; mobile phase A was 0.1 % formic acid in water; and mobile phase B was 0.1 % formic acid in acetonitrile. A/B gradient started at 5 % B for 3 min after injection and increased linearly to 95 % B at 13 min, then back to 5 % B over 0.1 min and finally held at 5 % B for an additional 1.9 min to re-equilibrate the column. Quantitation of indole was then achieved using standard curves generated using indole standard. Standard curve concentrations at 10, 50, 100, 300, 500, 700 and 9000 pmol/L in the solvent (methanol: water 20: 80 % (v/v)). The data were analyzed and processed using Xcalibur software (Thermofisher).

### Quantitative real-time RT-PCR analysis

Total RNA from worms was isolated using Trizol reagent (Invitrogen, Carlsbad, CA). Random-primed cDNAs were generated by reverse transcription of the total RNA samples using a standard protocol. A quantitative real-time-PCR analysis was performed with a Roche Light Cycler 480 System (Roche Applied Science, Penzberg, Germany) using SYBR Premix (Takara, Dalian, China). The relative amount of *lys-7* or *lys-8* mRNA to *act-1* mRNA (an internal control) was calculated using the method described previously (Pfaffl 2001). The primers used for PCR were as follows: *act-1*: 5’- CGT GTT CCC ATC CAT TGT CG -3’ (F), 5’- AAG GTG TGA TGC CAG ATC TTC -3’ (R); *lys-7*: 5’- GTC TCC AGA GCC AGA CAA TCC -3’ (F), 5’- CCA GTG ACT CCA CCG CTG TA -3’ (R); *lys-8*: 5’- GCT TCA GTC TCC GTC AAG GTC -3’(F), 5’- TGA AGC TGG CTC AAT GAA AC -3’ (R).

### Statistics

Differences in survival rates were analyzed using the log-rank test. Differences in mRNA levels, the percentage of body cavity-leakage, fluorescence intensity, and CFU were assessed by performing the unpaired t-test. Differences in DAF-16::GFP nuclear accumulation were analyzed using the two-way ANOVA. Differences in survival rates were analyzed using the log-rank test. Data were analyzed using GraphPad Prism 8.0.

## Acknowledgments

We thank the Caenorhabditis Genetics Center, Dr. Gary Ruvkun, and Dr. Mengqiu Dong for nematode strains. This work was supported by a grant from the National Key R&D Program of China (2022YFD1400700), the National Natural Science Foundation of China (U1802233 and 31900420), and the independent research fund of Yunnan Characteristic Plant Extraction Laboratory (2022YKZY006).

## Author Contributions

C.G.Z., Y.C.M., and R.Q.Y. designed the experiments and analyzed the data. R.Q.Y., Y.H.C., Q.Y.W., J.T., S.Z.N., and Q.Z performed the experiments. C.G.Z., Y.C.M., R.Q.Y., and Y.H. C. interpreted the data. C.G.Z. and Y.C.M. wrote the manuscript.

## Data availability statement

All data generated or analyzed during this study are included in the manuscript and supporting source data file.

---

**Figure 1-figure supplement 1.**
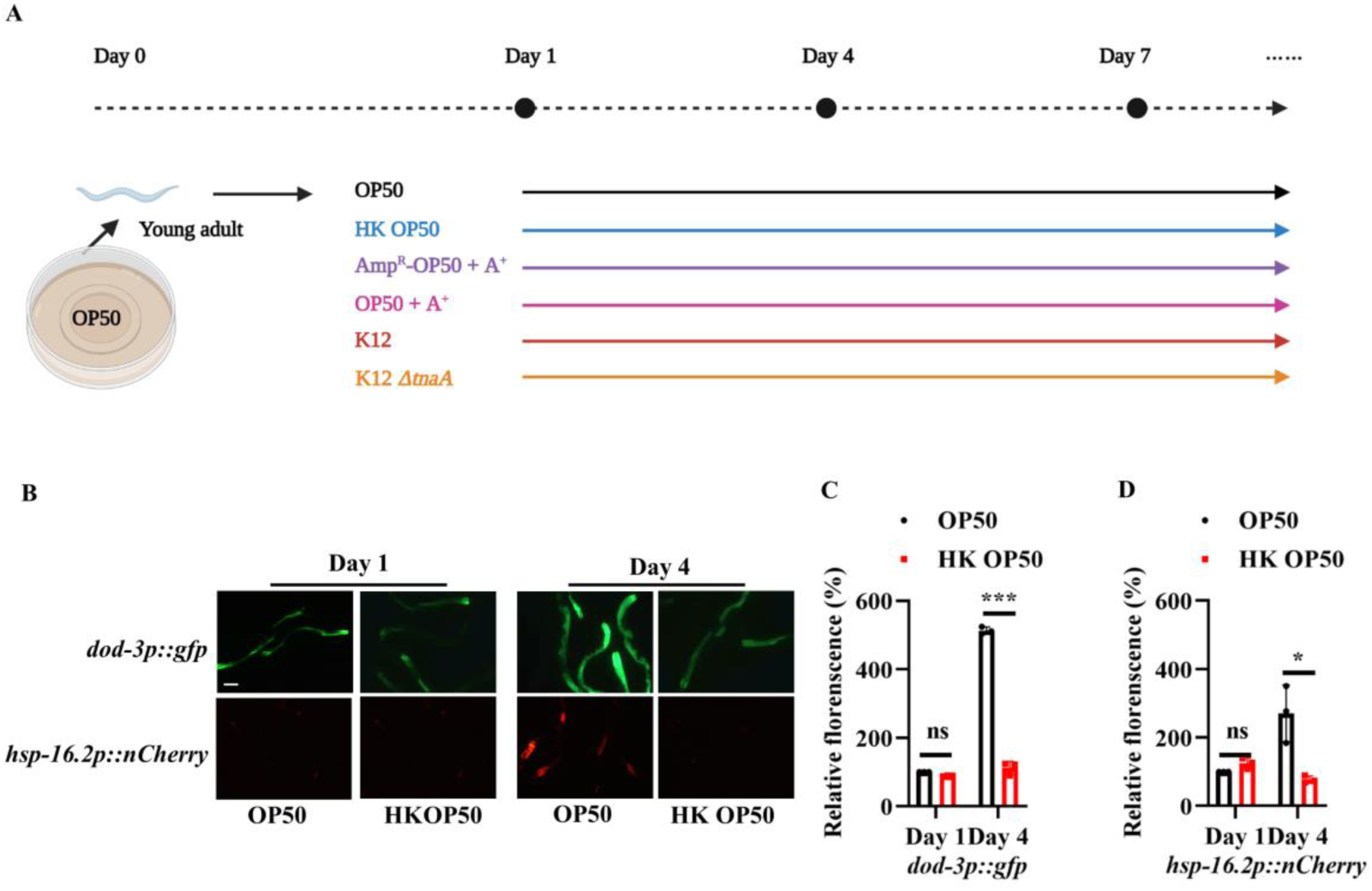
The expression of two DAF-16 target genes *dod-3* and *hsp-16.2* is significantly up-regulated in worms fed live *E. coli* OP50 on Day 4. (A) Study design for age-related dysbiosis study. (B) Representative images of worms expressing *dod-3p::gfp* and *hsp-16.2p::nCherry* fed live or heat-killed (HK) *E. coli* OP50 on Days 1 and 4. Scale bars: 100 μm. (C and D) Quantification of fluorescent intensity of *dod-3p::gfp* (C), and *hsp-16.2p::nCherry* (D). These results are means ± SEM of three independent experiments (n > 30 worms per experiment). \*\*\**P* < 0.001; \**P* < 0.05. ns, not significant. *P*-values (C and D) were calculated using the unpaired t-test. **Figure 1-figure supplement 1-source data 1** **Quantification results.**

**Figure 1-figure supplement 2.**
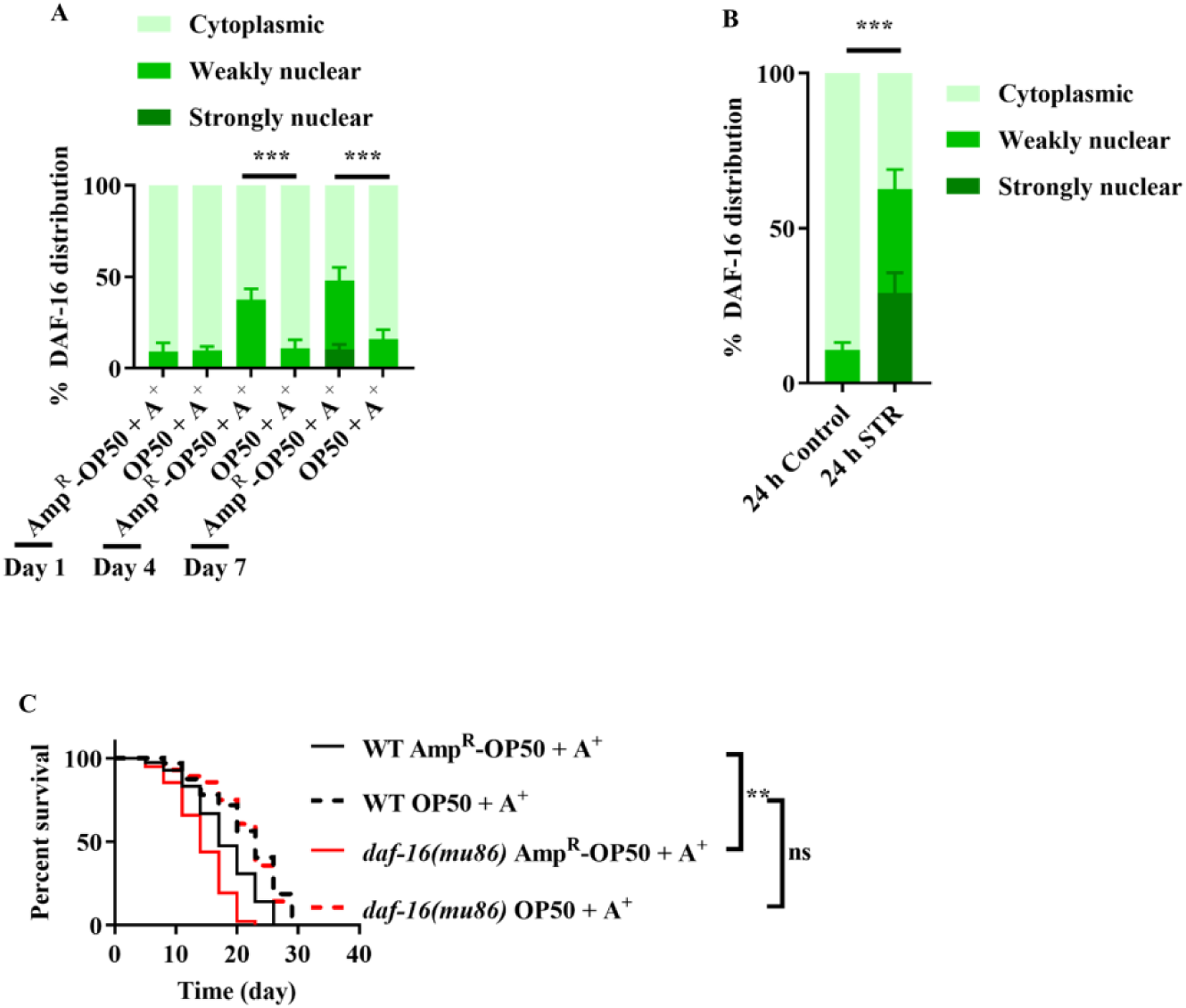
DAF -16 is not activated by antibiotic-killed *E. coli* OP50. DAF-16 was also retained in the cytoplasm of the intestine of worms fed ampicillin-killed *E. coli* OP50 on Days 4 and 7. These results are means ± SEM of three independent experiments (n > 35 worms per experiment). \*\*\**P* < 0.001. (B) Starvation induced the nuclear translocation of DAF-16 in the intestine of worms on Day 1. These results are means ± SEM of three independent experiments (n > 35 worms per experiment). \*\*\**P* < 0.001. *P*-values (**A** and **B**) were calculated using two-way ANOVA. (**C**) *daf-16(mu86)* mutants grown on live *E. coli* OP50 had a shorter lifespan compared to those grown on ampicillin-killed *E. coli* OP50. \*\**P* < 0.01. ns, not significant. A^+^, Ampicillin treatment; Amp^R^-OP50, *E. coli* OP50 containing an ampicillin resistance plasmid (PMF440). *P*-values throughout were calculated using log-rank test. **Figure 1-figure supplement 2-source data 1** **Lifespan assays summary and quantification results.**

**Figure 2-figure supplement 1.**
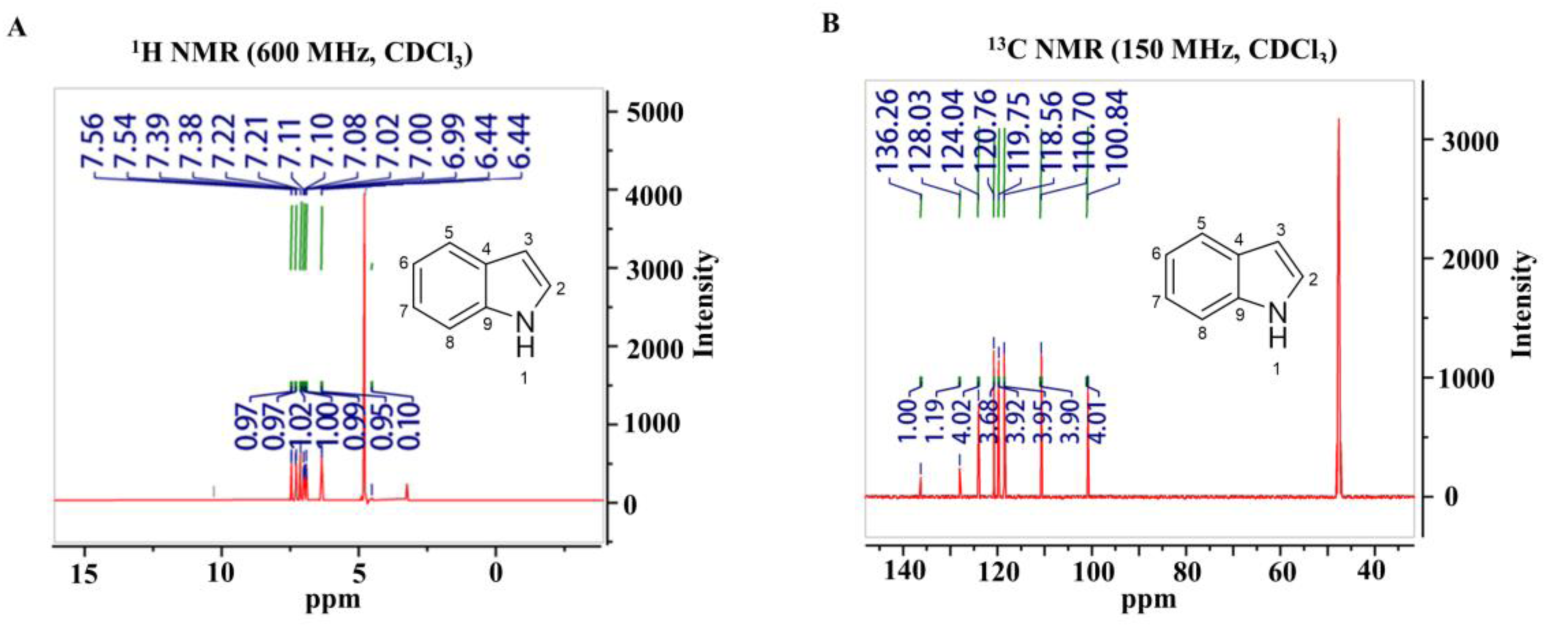
Indole is the active compound for activation of DAF-16. **(A and B)** The candidate compound for activation of DAF-16 was identified as indole with ^1^H NMR (A) and ^13^C NMR (B).

**Figure 2-figure supplement 2.**
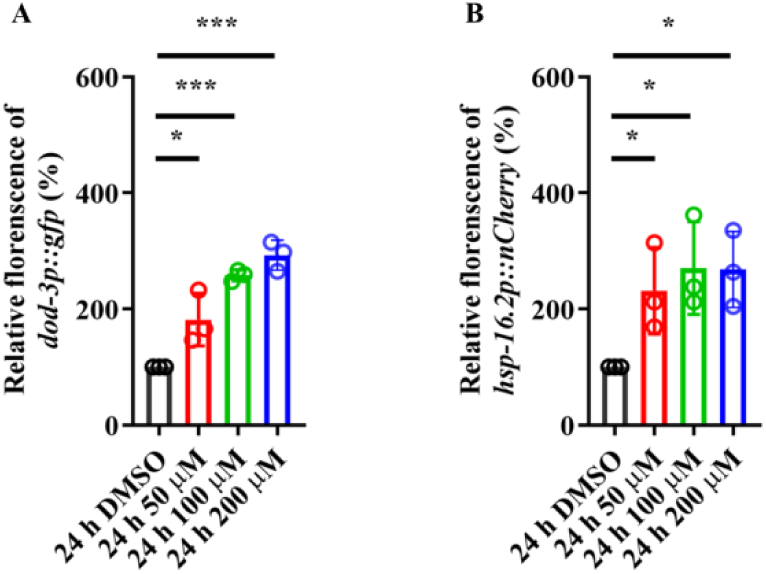
Indole treatment induces the expressions of two DAF-16 target genes in worms. **(A and B)** Quantification of fluorescent intensity of *dod-3p::gfp* (A) or *hsp-16.2p::nCherry* (B) in young adult worms treated with indole (100 μM). These results are means ± SEM of three independent experiments (n > 35 worms per experiment). \**P* < 0.05; \*\*\**P* < 0.001. *P*-values throughout were calculated using the unpaired t-test. **Figure 2-figure supplement 2-source data 1** **Quantification results.**

**Figure 2-figure supplement 3.**
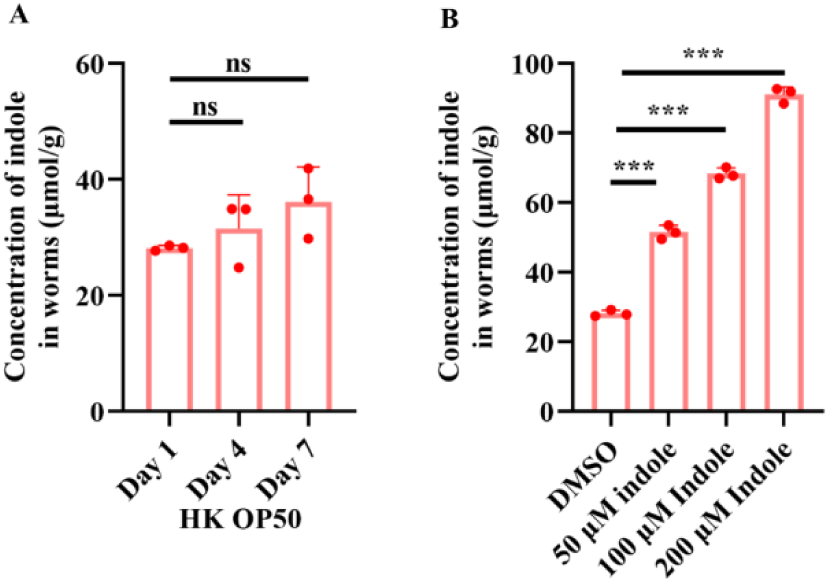
Quantitative analysis of indole in *C. elegans* by LC-MS. (**A**) The levels of indole were not altered in worms fed heat-killed (HK) *E. coli* OP50 on Days 1, 4, and 7. These results are means ± SEM of three independent experiments. ns, not significant. (**B**) Supplementation with indole (50-200 μM) significantly increased the indole levels in young adult worms after 24 h treatment. These results are means ± SEM of three independent experiments. \*\*\**P* < 0.001. *P*-values throughout were calculated using the unpaired t-test. **Figure 2-figure supplement 3-source data 1** **Quantification results.**

**Figure 2-figure supplement 4.**
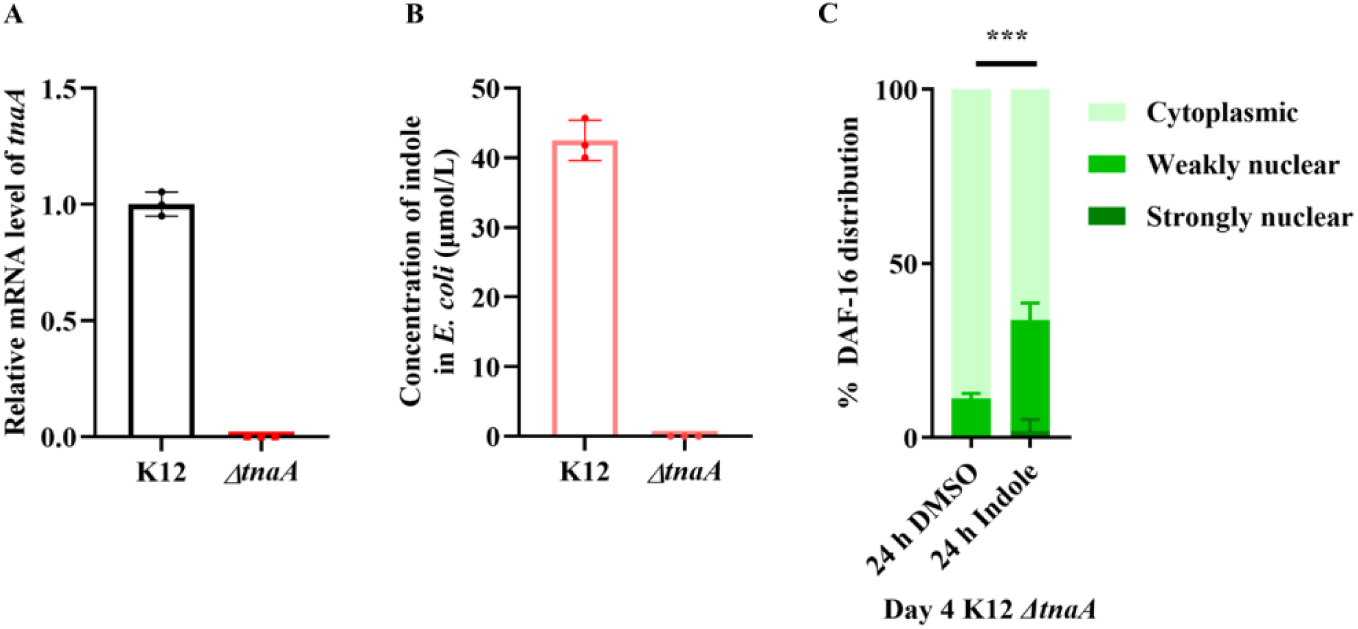
Functional validation of *tnaA*-deficient BW25113 strain. (**A and B**) The levels of *tnaA* mRNA (A) and indole (B) were undetectable in the K-12 *Δ tnaA* strains. These results are means ± SEM of three independent experiments. (**C**) The nuclear translocation of DAF-16::GFP was mainly located in the cytoplasm of the intestine in worms fed live K-12 *ΔtnaA* strains on Day 4. However, supplementation with indole (100 μM) induced the nuclear translocation of DAF-16::GFP in the intestine in these worms. These results are means ± SEM of three independent experiments. \*\**P* < 0.01. *P*-value was calculated using the Friedman test (with Dunn’s test for multiple comparisons). **Figure 2-figure supplement 4-source data 1** **Quantification results.**

**Figure 4-figure supplement 1.**
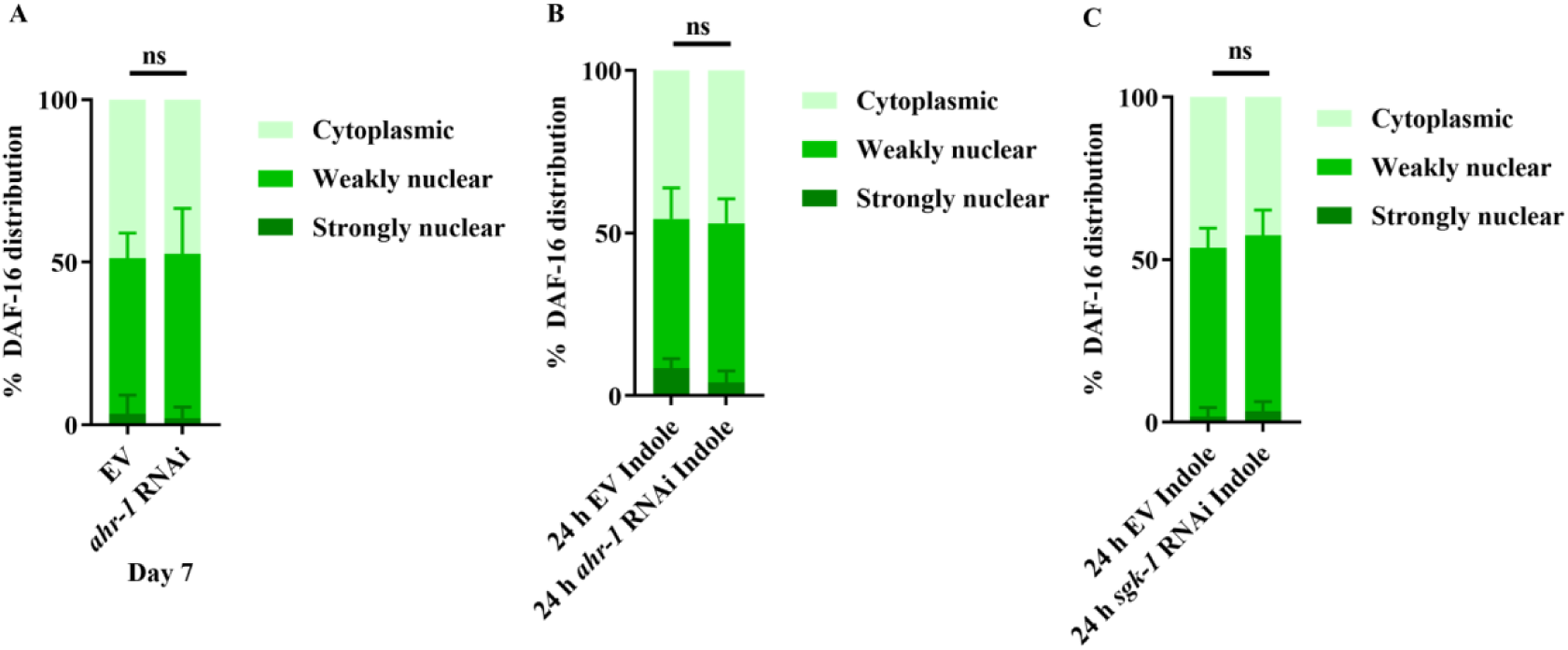
Indole promotes nuclear localization of DAF-16 independent of AHR-1 and SGK-1. **(A and B)** Knockdown of *ahr-1* by RNAi did not affect the nuclear translocation of DAF-16 in worms fed K12 strain on Day 7 (A) or young adult worms treated with indole for 24 h (B). These results are means ± SEM of three independent experiments (n > 35 worms per experiment). **(C)** Knockdown of *sgk-1* by RNAi did not affect the nuclear translocation of DAF-16 in young adult worms treated with indole (100 μM) for 24 h. These results are means ± SEM of three independent experiments (n > 35 worms per experiment). ns, not significant. *P*-value throughout was calculated using two-way ANOVA. **Figure 4-figure supplement 1-source data 1** **Quantification results.**

**Figure 4-figure supplement 2.**
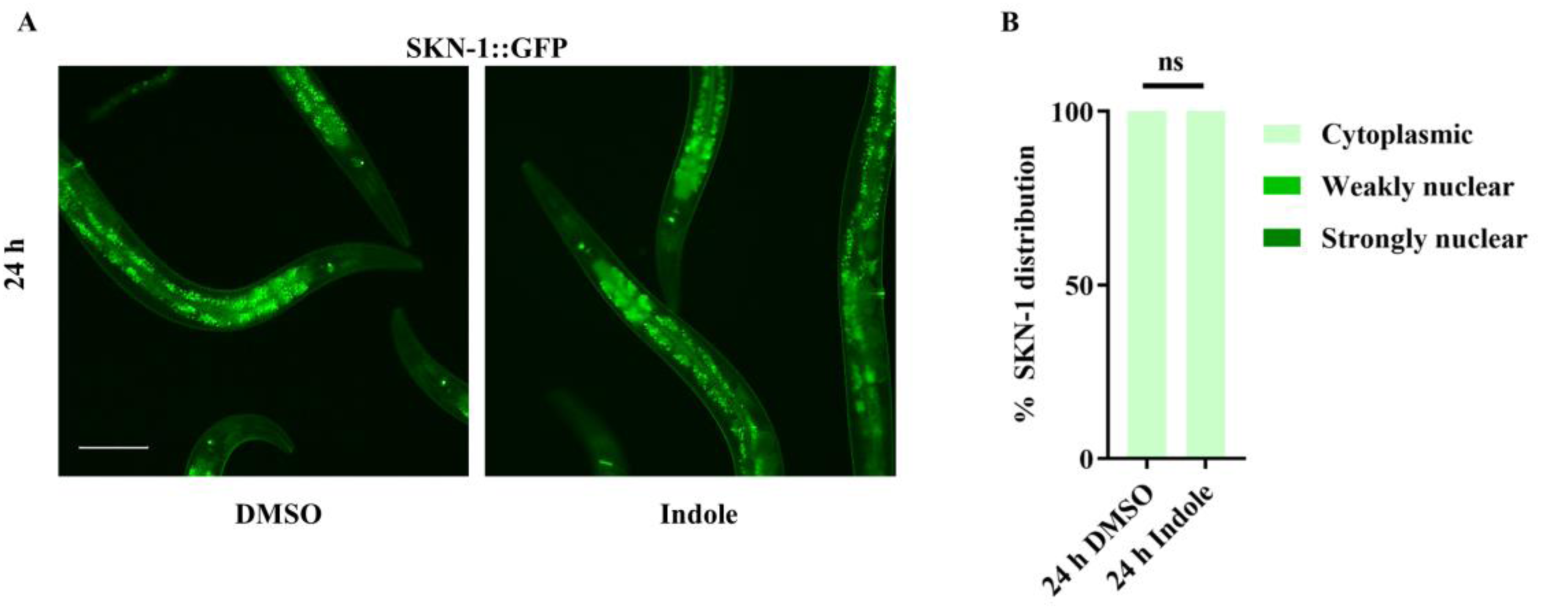
Supplementation with indole fails to induce nuclear localization of SKN-1::GFP in worms. (**A**) Representative images of SKN-1::GFP. Scale bars: 100 μm. (**B**) Quantification of SKN-1 nuclear localization. These results are means ± SEM of three independent experiments (n > 35 worms per experiment). ns, not significant. *P*-value was calculated using the using the two-way ANOVA. **Figure 4-figure supplement 2-source data 1** **Quantification results.**

**Figure 6-figure supplement 1.**
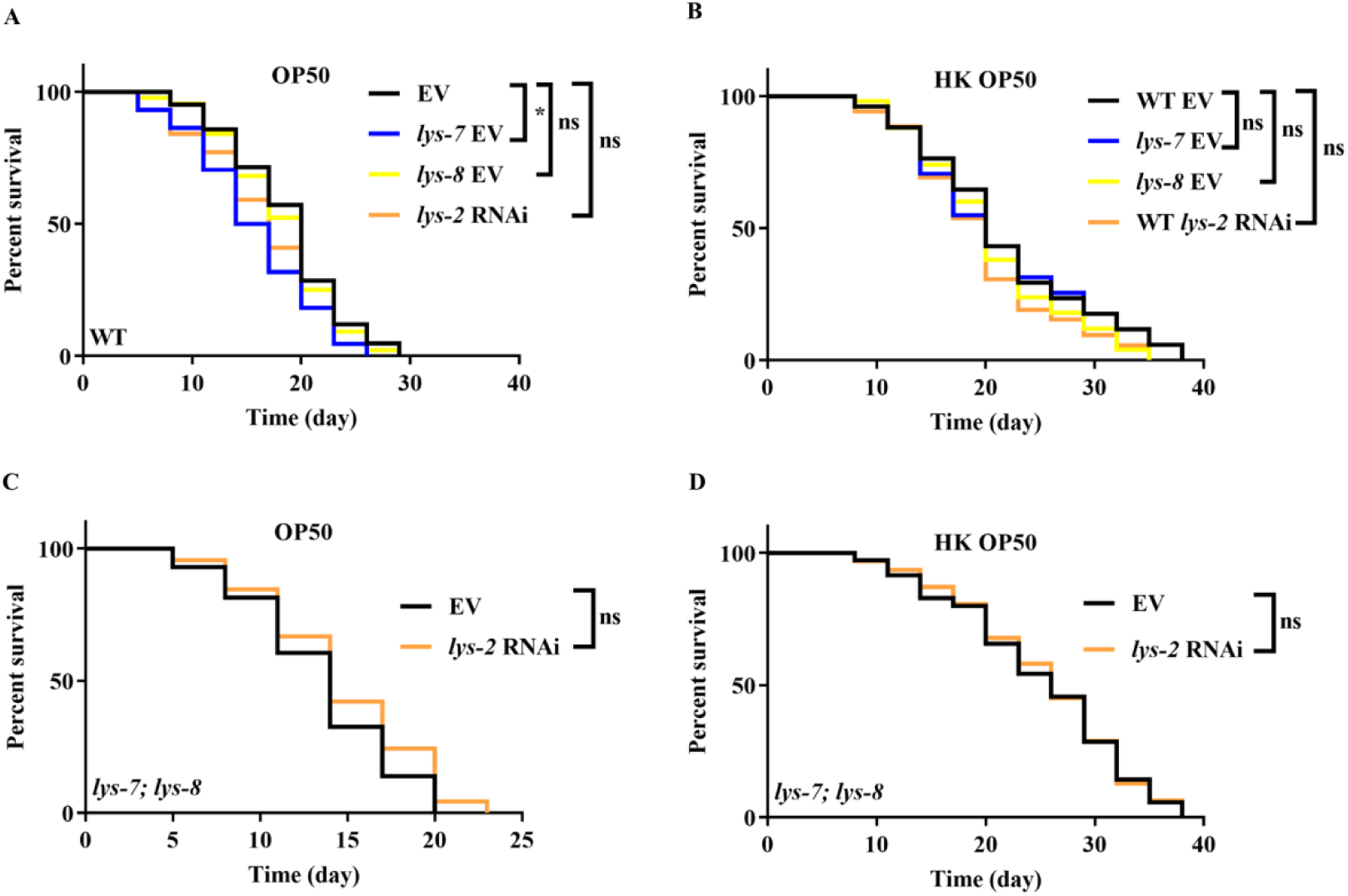
The roles of lysozyme genes in lifespan in worms. **(A and B)** The lifespans of *lys-7(ok1384)*, or *lys-8(ok3504)* mutants, or *lys-2* RNAi worms fed either live *E. coli* OP50 (A) or heat-killed (HK) *E. coli* OP50 (B). \**P* < 0.05. ns, not significant. **(C)** Knockdown of *lys-2* by RNAi did not affect the lifespan of *lys-7(ok1384)*; *lys-8(ok3504)* double mutants fed live (C) or HK *E. coli* OP50 (D). ns, not significant. *P*-values throughout were calculated using log-rank test. **Figure 6-figure supplement 1-source data 1** **Lifespan assays summary.**

**Figure 6-figure supplement 2.**
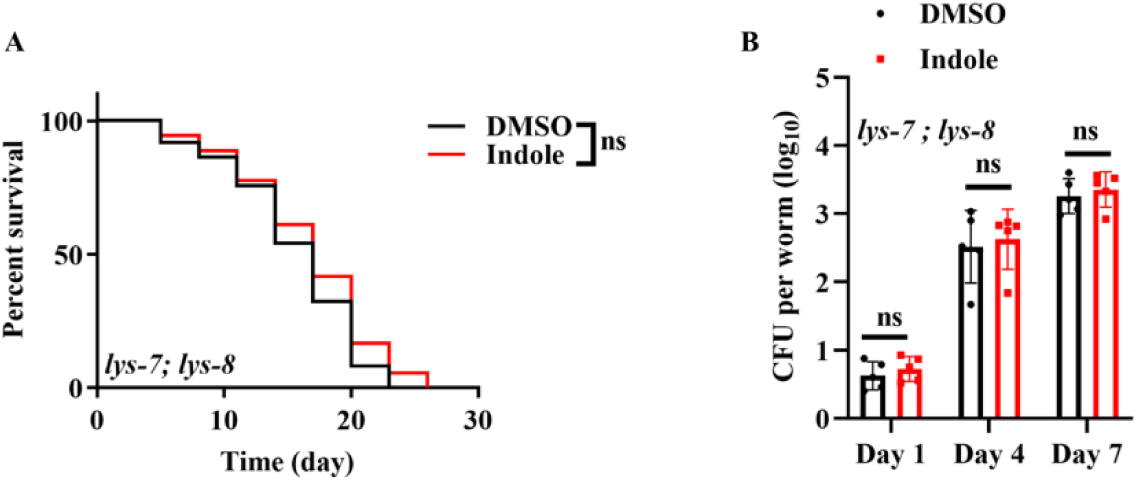
Indole-mediated lifespan extension in worms depends on LYS-7 and LYS-8. (**A**) Supplementation with indole (100 μM) no longer extended lifespan in *lys-7(ok1384); lys-8(ok3504)* double mutants. *P*-values throughout were calculated using log-rank test. (**B**) Supplementation with indole failed to suppress the increase the CFU of K-12 in *lys-7(ok1384); lys-8(ok3504)* double mutants. These results are means ± SEM of five independent experiments (n > 30 worms per experiment). ns, not significant. *P*-values were calculated using the unpaired t-test. **Figure 6-figure supplement 2-source data 1** **Lifespan assays summary and quantification results.**

**Figure 7-figure supplement 1.**
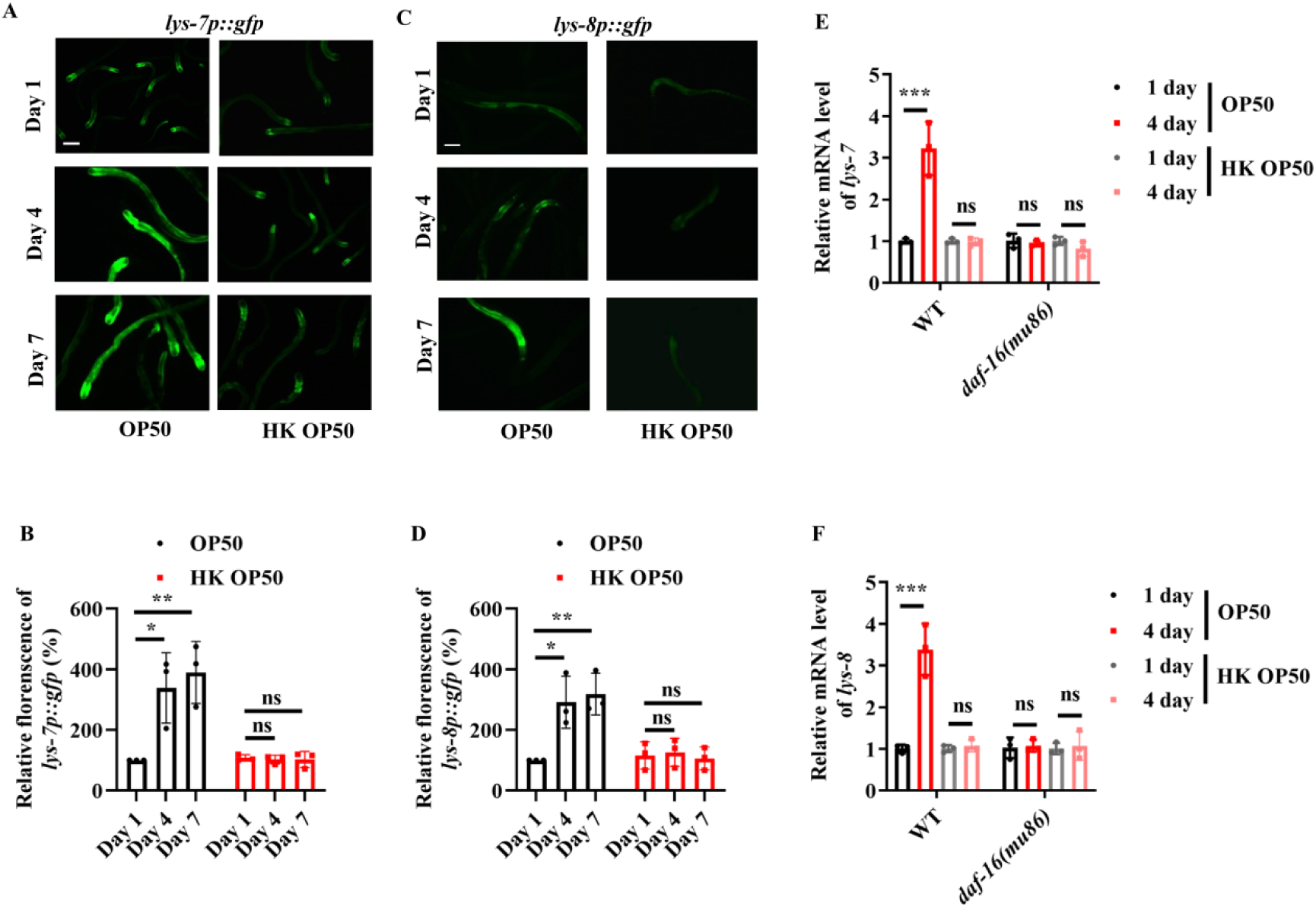
The expressions of *lys-7p::gfp* and *lys-8p::gfp* are up-regulated in worms with age. **(A and C)** Representative images of *lys-7p::gfp* (A) and *lys-8p::gfp* (C) in worms fed live *E. coli* OP50. Scale bars: 100 μm. **(B and D)** Quantification of fluorescent intensity of *lys-7p::gfp* (B) and *lys-8p::gfp* (D). The expressions of *lys-7p::gfp* and *lys-8p::gfp* were significantly up-regulated in worms fed live *E. coli* OP50, but not heat-killed (HK) *E. coli* OP50, on Days 4 and 7. These results are means ± SEM of three independent experiments (n > 35 worms per experiment). \**P* < 0.05; \*\**P* < 0.01. **(E and F)** The mRNA levels of *lys-7* (E) and *lys-8* (F) are up-regulated in wild-type (WT) worms fed live *E. coli* OP50, but not HK *E. coli* OP50, on Day 4. These increases in the mRNA levels of *lys-7* (E) and *lys-8* (F) were abolished by a mutation in *daf-16(mu86)*. \*\*\**P* < 0.001. These results are means ± SEM of three independent experiments. *P*-values (**B,** and **D-F**) were calculated using the unpaired t-test. **Figure 7-figure supplement 1-source data 1** **Quantification results.**

**Figure 7-figure supplement 2.**
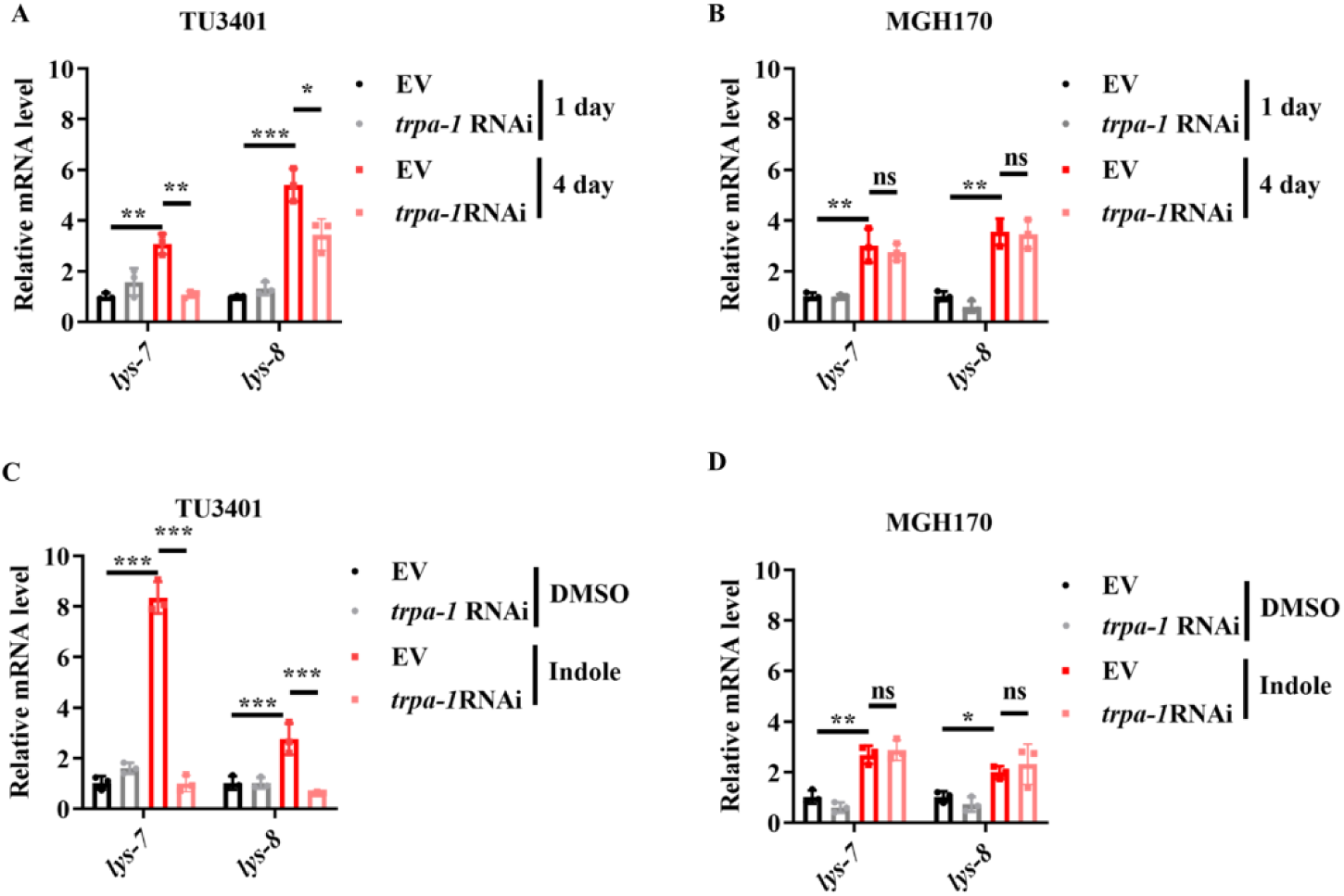
TRPA-1 in neurons is required for the expression *lys-7* and *lys-8*. (**A and B**) The mRNA levels of *lys-7* and *lys-8* were significantly down-regulated in worms subjected to neuronal-specific (A), but not intestinal-specific (B), knockdown of *trpa-1* by RNAi. These results are means ± SEM of three independent experiments. \*\*\**P* < 0.001. \*\**P* < 0.01. \**P* < 0.05. ns, not significant. (**C and D**) Supplementation with indole (100 μM) up-regulated the mRNA levels of *lys-7* and *lys-8* in worms subjected to neuronal -specific (C), but not intestinal -specific (D), knockdown of *trpa-1* by RNAi. These results are means ± SEM of three independent experiments. \*\**P* < 0.01; \*\*\**P* < 0.001. ns, not significant. *P*-values (**A-D**) were calculated using the unpaired t-test. **Figure 7-figure supplement 2-source data 1** **Quantification results.**

**Table S1.** The ^1^H and ^13^C NMR spectroscopic data of indole at 600 MHz for ^1^H NMR and 150 MHz for ^13^C NMR with reference to the solvent signals. NMR spectra of indole were recorded in CDCl_3_. The corresponding ^1^H and^13^C NMR spectra were depicted in **Figure 2-figure supplement 1**, respectively. δ_H_ were recorded at 600 MHz and the measured values of δ_H_ were in good agree with published NMR data for indole. δ_C_ were recorded at 150 MHz and the values exhibited a good consistency with published data in ppm (Yagudaev, 1986).

| Atom position | $^1\text{H}$ NMR ($\delta_{\text{H}}$) | $^{13}\text{C}$ NMR ($\delta_{\text{C}}$) |
| --- | --- | --- |
| 1 | - | - |
| 2 | 7.22 (1H, d, $J = 3.0$ Hz) | 124.0 |
| 3 | 6.44 (1H, d, $J = 3.0$ Hz) | 100.9 |
| 4 | - | 128.0 |
| 5 | 7.55 (1H, d, $J = 7.8$ Hz) | 119.8 |
| 6 | 7.00 (1H, t, $J = 7.8$ Hz) | 120.8 |
| 7 | 7.10 (1H, t, $J = 7.8$ Hz) | 118.6 |
| 8 | 7.38 (1H, d, $J = 7.8$ Hz) | 110.7 |
| 9 | - | 136.3 |

